# An Eco-Evolutionary Framework to Predict Long-term Strain Diversity and Turnover in Infectious Diseases

**DOI:** 10.1101/2025.08.08.669409

**Authors:** Jiawei Liu, Qixin He

## Abstract

Pathogen strain diversity varies widely across diseases, yet the processes that govern its emergence, maintenance, and turnover remain unclear. We present MultiSEED, a fast numerical framework that captures strain dynamics across ecological and evolutionary timescales. Unlike previous models, MultiSEED distinguishes between transient and established strains, allowing classification of diseases into regimes of extinction, replacement, or coexistence. When applied to influenza A, HIV-1, pneumococcus, and malaria, the model reproduces observed strain diversity and turnover while revealing the key drivers of these differences. We show that susceptible host replenishment is central to long-term strain persistence, host population size and *R*_0_ shape overall diversity, and innovation rates strongly influence dynamic regimes, with their combined interactions ultimately determining pathogen strain diversity and turnover. This unifying framework can be readily applied across systems, enabling the rapid prediction of pathogen responses to environmental and demographic changes and guiding strategies for disease control.

---

The evolution of lymphocytes with diverse antigen receptors marked a key innovation in vertebrate immunity, enabling recognition, memory, and targeted elimination of pathogens (*1*). In jawless and jawed vertebrates, this diversity arises through recombination of leucine-rich repeat sequences or V(D)J gene segments, respectively, generating large repertoires capable of detecting a vast range of antigens with high precision (*2*). In response, pathogens have developed a counter-strategy: antigenic variation, a process that continually alters their surface proteins or polysaccharides to evade immune detection (*3*). This strategy hinders the development of sterilizing immunity, allowing for persistent or recurrent infection (*4, 5*). Given its success, one might expect antigenic variation to be widespread across pathogens, driving high strain diversity in most systems. Yet, this is not what we observe.

Pathogen strain diversity varies widely among diseases, often in ways that defy intuitive expectations. Some pathogens, such as *Streptococcus pneumoniae* and *Neisseria* spp., maintain extensive antigenic diversity despite moderate transmissibility (*3, 6*). In contrast, viruses, such as measles and chickenpox, are highly transmissible and mutate rapidly, yet circulate as single dominant strains (*7*). The potential tradeoff between pathogen transmissibility and strain diversity (*8*) raises a fundamental question in disease ecology: what governs the emergence, maintenance, or collapse of strain diversity in host-pathogen systems?

Previous modeling efforts have shown that higher basic reproduction numbers (*R*_0_) and longer infection durations relative to host lifespan promote higher diversity (*9–11*), while low mutation rates (*12, 13*) or transient cross-immunity (*14*) are essential for the low diversity and rapid replacement of dominant strains observed, as seen in influenza A. Additionally, a more structured host contact network (*8, 15*) and higher host population sizes (*9*) have been shown to increase strain diversity. Yet, we lack a general theoretical framework that can quantitatively predict the diversity and turnover dynamics of pathogen strains across both ecological and evolutionary timescales.

Agent-based stochastic models have been designed to track the long-term strain dynamics (*12, 13, 16*), but they focus on specific pathogens or isolate the effects of individual parameters, making it difficult to predict strain dynamics under the combined effect of key disease parameters. Conversely, equation-based deterministic models typically lack the dynamic emergence of *de novo* strains (summarized in (*17*)). They often assume a fixed set of strains and are primarily used to study equilibrium strain structures (*18*) or predict the invasibility of a new strain (*19*). While valuable for theoretical insights, these models frequently diverge from the outcomes observed in stochastic simulations (*11, 20*). Invasibility analyses tend to overestimate the success of new strains when the frequent extinctions that occur during epidemic troughs (termed overcompensatory effect) are not accounted for (*19*). These discrepancies highlight the critical role of ecological-timescale stochasticity in shaping long-term evolutionary patterns of strain dynamics (*21, 22*).

A more integrative effort by Abu-Raddad & Ferguson (*9*) linked cross-immunity and mutation to long-term diversity using an analytical framework (hereafter AF-model). Although this approach considered the strain death rate on the evolutionary timescale, it did not incorporate extinction during the early epidemic troughs on the ecological timescale, potentially leading to an overestimated invasion success of new strains. In addition, without characterizing the contribution of transient and established strains to disease dynamics, it is unable to explain why some pathogens are dominated by long-lasting endemic strains while others experience constant epidemic renewal.

Moreover, the impact of two important biological factors on strain dynamics remains poorly understood: finite specific immunity and within-host mixed infection. Many infectious diseases do not elicit lifelong immunity to a given strain, with immune protection waning over time (*5, 23–26*). Part of the problem is that this effect of immunity loss has rarely been incorporated into multi-strain modeling frameworks, despite its potential importance for long-term pathogen dynamics (but see (*19*)). Similarly, mixed infections, i.e., simultaneous infections by multiple antigenically distinct strains, have been observed in over 50 human diseases and many animal diseases (*27, 28*). The prevalence of mixed infections is influenced by transmission intensity, clinical risk (*29*), host immunity, and host age (*30*). Importantly, within-host competition resulting from mixed infections (*28*) and its impact on transmissibility and strain persistence have not been captured by multi-strain models.

Together, these limitations impede efforts to explain the vastly different strain diversity and dynamics between pathogens. We argue that what is needed is a general, analytically tractable computational framework that accounts for demographic stochasticity across ecological and evolutionary time scales, while integrating the waning of specific immunity and mixed infections. We address this gap with **<u>M</u>**ulti-**<u>S</u>**train **<u>E</u>**co-**<u>E</u>**vo **<u>D</u>**ynamics (MultiSEED), a theoretical framework that unifies epidemiological, immunological, and evolutionary processes in multistrain pathogens. This framework enables analytical predictions of both strain diversity and turnover dynamics, including the number of circulating strains, the turnover rates of emergent versus established variants, and the conditions under which strain coexistence or replacement dominates. We validate the numerical solutions from MultiSEED using individual-based simulations and we compare its predictions to observed diversity patterns in influenza A, HIV-1, *Streptococcus pneumoniae* (hereafter pneumococcus), and *Plasmodium falciparum* (hereafter malaria). Our framework not only addresses key theoretical questions, such as why some pathogens maintain stable diversity while others do not, but also supports practical applications, including predictions of vaccine update frequency, estimates of the risk of mixed infections, and recommendations as to whether intervention strategies should target transmission, reservoirs, or vector control.

## MultiSEED framework

To model the transmission dynamics and long-term persistence of multiple co-circulating strains, MultiSEED integrates a deterministic *n*-strain SIRS (Susceptible-Infected-Recovered-Susceptible) model with continuous-time stochastic processes at both ecological and evolutionary timescales (Fig. 1, details in SM, Fig. S1). The deterministic component extends the classic SIRS model to a multi-strain, status-based formulation, which does not require a record of individual infection histories, thus vastly reducing the number of equations (Fig. 1A, Fig. S1A). Instead, it adopts a pathogen-centric perspective, tracking host immune states and current infection states with respect to each strain (see also (*31*) and (*32*)). We incorporate two forms of immunity status that reflect key features of multi-strain pathogen dynamics: (1) strain-specific immunity, which prevents reinfection by the same strain but wanes over time, and (2) lifelong cross-immunity, which reduces susceptibility to other strains following any infection.

**Figure 1.**
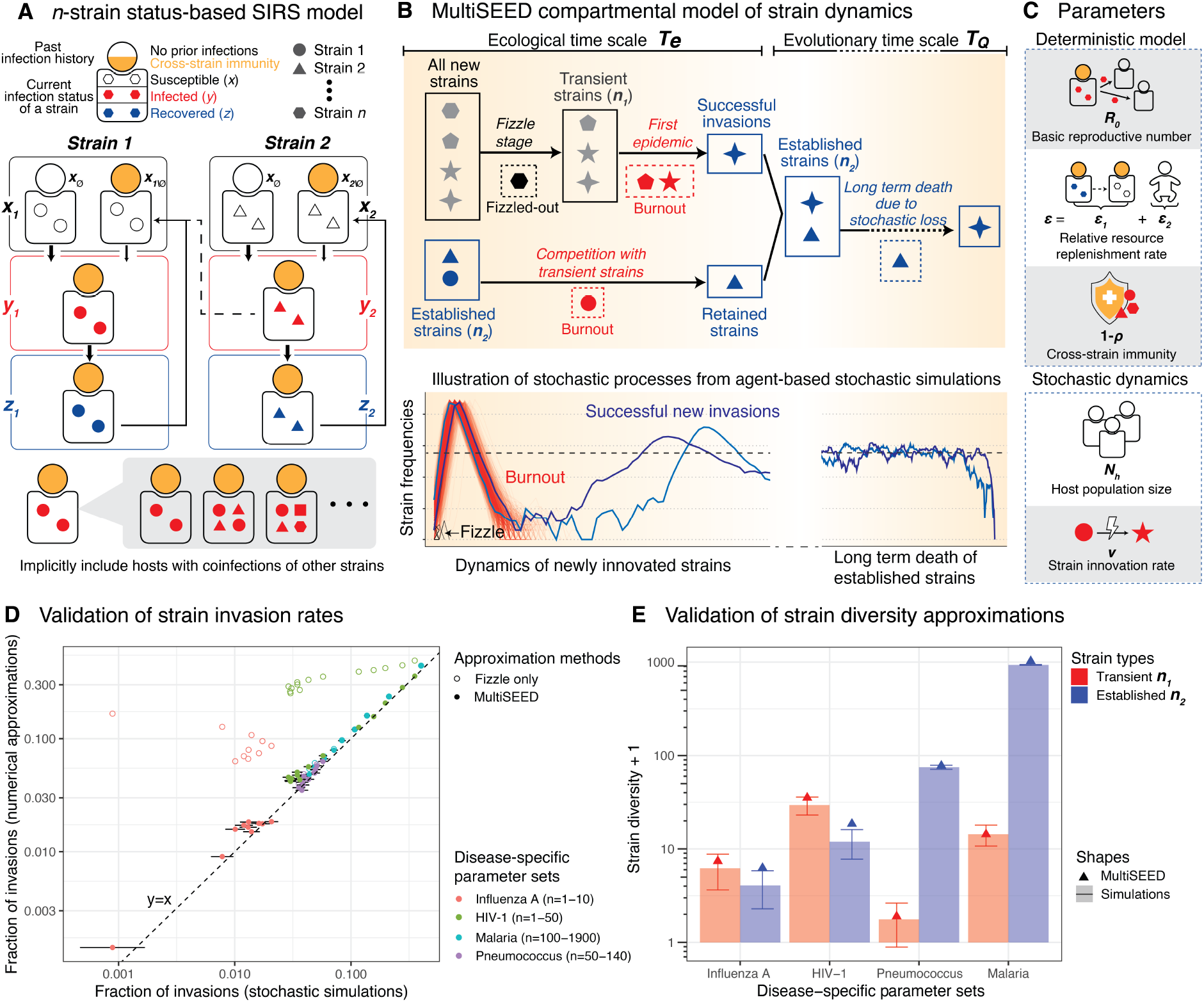
MultiSEED framework and the validation results. **(A)** The deterministic *n*-strain status-based SIRS model. The classes of susceptible, infected and recovered host proportions for strain *i* are referred as *x*_*i*_, *y*_*i*_ and *z*_*i*_. Hosts infected with strain *i* may simultaneously carry infections from other strains. **(B)** An illustration of continuous time stochastic processes in strain dynamics over the ecological (left) and evolutionary (right) time scales. Strains are categorized into transient (*n*_1_, gray) and established (*n*_2_, blue) groups. Dashed boxes indicate strain extinctions. Refer to Section S1.5 of the SM for the compartmental ODE model describing the strain dynamics. **(C)** The model parameters as factors affecting strain diversity and regimes of strain dynamics. **(D-E)** Validation of MultiSEED by comparison with the simulation results from an agent-based model, under parameter sets specific to each disease: Influenza A: *ε* = 0.005, *ε*_1_ = 0.0045, *R*_0_ = 2; HIV-1: *ε* = 0.022, *ε*_1_ = 0.002, *R*_0_ = 3; Pneumococcus: *ε* = 0.013, *ε*_1_ = 0.0065, *R*_0_ = 1.5; Malaria: *ε* = 0.035, *ε*_1_ = 0.026, *R*_0_ = 5. Other parameters can be found in Table 1. **(D)** Comparison of the invasion probability of new strains into a system with *n* resident strains between numerical approximations and stochastic simulations; error bars represent 95% binomial proportion confidence interval. **(E)** Comparison of the number of transient and established strains at equilibrium between numerical approximations and stochastic simulations; error bars represent standard deviations of simulation runs.

In this study, we define strains as immunologically distinct phenotypes, rather than tracking fine-scale mutational pathways or the genetic architecture underlying antigenic diversification (*33, 34*). Prior work has shown that immune selection tends to organize pathogen populations into discrete, minimally overlapping antigenic clusters (*16, 32, 34*). Building on this insight, we thus focus on predicting the effective number of distinct immunological entities circulating in the host population over time, providing a coarse-grained, population-level perspective that complements mutation-level or allele-based analyses.

**Table 1:** Estimated parameter values for the four focal human diseases from literature: influenza A, HIV-1, pneumococcus and malaria, along with corresponding literature sources in brackets.

| Disease | $N_h$ | $\nu$ | $\varepsilon$ | $R_0$ | $\rho$ | #Gens/year | #Years |
| --- | --- | --- | --- | --- | --- | --- | --- |
| Influenza A | $10^6$ (55) | $10^{-4}$ (38, 56–61) | $0.003 - 0.006$ (26, 59–61) | $1.2 - 1.85$ (62) | $0.5$ (63) | $52$ (59–61) | $1000$ (64) |
| HIV-1 | $10^4$ (65) | $10^{-3}$ (66–73) | $0.013 - 0.038$ (66, 74–76) | $1.6 - 3.1$ (77) | $0.5$ (78) | $1$ (66) | $100$ (39, 40) |
| Pneumococcus | $10^6$ (79) | $10^{-6}$ (80–85) | $0.0065 - 0.023$ (80, 81, 86–88) | $1.23 - 2$ (89–91) | $0.7$ (92–95) | $3$ (80, 81) | $100000$ (85) |
| Malaria | $10^5$ (96) | $10^{-5}$ (16, 97–100) | $0.033 - 0.059$ (23, 97–99, 101–103) | $\geq 1$ (104) | $0.7$ (105) | $2.5$ (97–99) | $10000$ (106, 107) |

For each strain *i*, hosts are categorized into three compartments: the fractions of susceptible (*x*_*i*_), infected (*y*_*i*_), and recovered (*z*_*i*_). The susceptible class *x*_*i*_ is further subdivided into fully naïve hosts (*x*_∅_) and experienced hosts with reduced susceptibility due to prior exposure to other strains or waned specific immunity to strain *i* (*x*_*i\*∅_). This formulation also allows for the convenient incorporation of mixed infections, where a single host can harbor multiple genetically distinct strains of the same pathogen species (Fig. 1A). As a result, the sum of *y*_*i*_ across strains can exceed 1. Within-host interactions among co-infecting strains take the form of competition for host resources, which reduces per-strain pathogen density and thereby decreases transmissibility. We denote this competition-induced reduction in baseline transmissibility *β* by *β*^*^, and the corresponding reduced reproduction number as 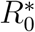. At the between-host level, co-circulating strains accelerate transitions from naïve hosts to experienced hosts (Fig. 1A), thereby reducing the hosts’ overall susceptibility to each strain. The infection recovery rate (*γ*) and the rate of strain-specific immunity loss (*δ*) are assumed to be unaffected by inter-strain competition. The host population size is held constant at demographic equilibrium, with the birth rate equal to the natural death rate (*µ*), and infection does not contribute to mortality. For analytical and computational tractability, we assume all circulating strains are epidemiologically equivalent and differ only in their antigenic space. This symmetry enables us to derive a single set of equations that characterizes the equilibrium behavior representative of all strains, along with an additional set describing the invasion dynamics of a novel strain introduced into a resident strain population (Eq. S9 and Eq. S10 in SM).

After rescaling time in the model by the expected duration of infection (i.e., 1*/*(*γ* + *µ*)), the dynamics of each strain are governed by three key parameters: the basic reproduction number (*R*_0_), the relative resource replenishment rate (*ε*), and the cross-strain susceptibility (*ρ*) (Fig. 1C). The parameter *ε* = *ε*_1_ + *ε*_2_ captures the rate at which the pool of strain-specific susceptible hosts is replenished relative to the pathogen’s lifespan. This replenishment occurs through two mechanisms: waning of strain-specific immune memory (*ε*_1_ = *δ/*(*γ* + *µ*)) and the birth of immunologically naïve individuals (*ε*_2_ = *µ/*(*γ*+*µ*)). For a detailed derivation and interpretation of *ε*, see SM section S1.1 and S3.1.

To capture how new strains invade and how they increase the chance of resident strains dying out, we formulate two classes of stochastic processes—one operating on the ecological timescale and the other on the evolutionary timescale (Fig. 1B). New strains with adequate antigenic distance from existing strains emerge continuously through intrinsic genetic processes, such as mutation. Thus, the total rate of new innovations is proportional to the host population size *N*_*h*_, the strain innovation rate *ν*, and the disease prevalence. However, not all novel strains successfully establish; they must overcome two key stochastic bottlenecks to persist within the system.

The first barrier, which we term the fizzle phase, occurs when a newly formed strain fails to generate enough secondary infections and dies out shortly after introduction (*35*). If a strain escapes the fizzle phase, it proceeds to its first epidemic wave. However, this expansion is followed by a sharp depletion of susceptible hosts towards the end of an epidemic, which can lead to a second extinction event, referred to as epidemic burnout (*22*) (Fig. 1B). During this invasion process, resident strains may also face stochastic extinction due to reductions in their infected host populations (SM Section S1.3; Fig. S2).

Strains that survive both the fizzle phase and epidemic burnout are considered successful invaders and transition into long-term circulation. While they may still experience stochastic fluctuations during subsequent oscillatory dynamics, the magnitude of these effects is considerably smaller (*22*). Extending the approach of Parsons et al. (*22*) for single-strain models to the multi-strain case, we numerically approximate the probabilities of fizzle and burnout using a combination of deterministic dynamics and time-inhomogeneous stochastic birth–death processes (see SM Section S1.3). For surviving strains that reach endemic levels, we assume quasi-stationary distributions of infected and susceptible hosts for the specific strain. The longterm extinction dynamics of these established strains can then be approximated by an Ornstein–Uhlenbeck (OU) process (*36, 37*) (SM Section S1.4).

Considering both the short-term invasion dynamics and the long-term extinction rates of strains, we develop a compartmental model to track strain turnover and predict long-term strain diversity (Fig. 1B, Fig. S1B). Strains are categorized into two classes: transient strains (*n*_1_), which have survived the fizzle phase but not yet completed their first epidemic, and established strains (*n*_2_), which have completed their first epidemic (Fig. 1B). When the rates of strain invasion and extinction are balanced, the total number of co-circulating strains at equilibrium is given by *n* = *n*_1_ + *n*_2_ (SM Section S1.5; Eq. S32).

The structure of this model allows us to delineate three distinct regimes of strain dynamics based on the relative abundance and lifespan of transient and established strains at equilibrium: 1) Extinction, where no strain can persist over evolutionary timescales; 2) Replacement, where transient and established strains occur at similar frequencies and lifespans, indicating continual turnover; 3) Coexistence, where established strains dominate in both abundance and longevity. To differentiate between the replacement and coexistence regimes, we use two criteria: 1) the proportion of transient strains, *n*_1_*/*(*n*_1_ +*n*_2_), and 2) the ratio between the epidemic burnout time and the expected persistence time of strains after establishment, *T*_*e*_*/*(*T*_*e*_ + *T*_*Q*_). Low values for both metrics signal a long-term coexistence regime. We evaluated cutoff thresholds from 0.1 to 0.4 to delineate these regimes and present results using a threshold of 1/3 in the main text (see alternative cutoffs in Fig. S5).

### Validation of MultiSEED using an agent-based simulator

We first evaluated whether MultiSEED accurately estimates the invasion probability of a novel strain in the presence of *n* resident strains by comparing its numerical approximations to results from an agent-based simulation model (adapted from (*16*); see SM Section S1.7). To assess performance across a broad range of epidemiological scenarios, we tested representative parameter sets for four major human pathogens: influenza A, HIV-1, pneumococcus, and malaria, spanning a wide range of assumed resident strain numbers. When both the fizzle phase and epidemic burnout are incorporated as prerequisites for successful invasion, the numerically approximated probabilities of invasion closely match those obtained from stochastic simulations (filled circles in Fig. 1D). In contrast, when only the fizzle phase is considered, an approach taken in previous studies (e.g., AF-model in (*9*)), the numerical approximations consistently overestimate invasion probabilities, especially for influenza A and HIV-1 (open circles in Fig. 1D). This discrepancy reflects the substantial impact of epidemic burnout on these diseases (Fig. S3).

We then assessed the quality of numerical approximations for long-term strain diversity by MultiSEED (Fig. 1E). The model predictions for the number of transient and established strains at equilibrium generally fall within the range of outcomes observed in stochastic simulations, particularly for transient strains, demonstrating strong agreement. However, for the parameter regimes of influenza A and HIV-1, MultiSEED overestimates the number of established strains by approximately a factor of two relative to simulation means. This overestimation is primarily due to the known difficulty that the Ornstein–Uhlenbeck (OU) approximation has in estimating the expected lifespan of an endemic strain (*T*_*Q*_) (*36*). The OU approximation of *T*_*Q*_ is thus less accurate in diseases for which many strains stochastically go extinct during successive epidemic troughs before reaching the quasi-stationary endemic state assumed by the OU model.

Despite these deviations for some extreme parameter regimes, the predicted invasion probabilities and strain diversity estimates from MultiSEED generally show strong agreement with results from full agent-based simulations. Thus, MultiSEED provides a useful framework for estimating long-term strain diversity and turnover to the correct order, even in systems characterized by complex or volatile dynamics.

### Estimating strain diversity and regimes of strain dynamics in focal human diseases

To validate MultiSEED’s ability to capture realistic strain dynamics, we assessed its predictions of strain composition, lifespan distributions, and turnover regimes in four human infectious diseases with well-documented variation in strain diversity and strain dynamics: influenza A, HIV-1, pneumococcus, and malaria (see Table 1 for parameter ranges and SM Section S1.8 for data sources and derivation methods). Among the five model parameters (Fig. 1C), the basic reproduction number *R*_0_ and the relative resource replenishment rate *ε* strongly shape transmission dynamics, yet their empirical estimates remain uncertain. We therefore explored a range of (*ε, R*_0_) combinations for each disease while keeping the other parameters (*N*_*h*_, *ν, ρ*) fixed.

The four focal diseases exhibit drastically different predicted strain diversity and the corresponding strain compositions (transient or established) within their empirically plausible parameter ranges (Fig. 2A and D, Fig. S4A). For influenza A, MultiSEED predicts either disease extinction or circulation of a small number of strains (between 1 and 10) at higher values of *ε* and *R*_0_, with both transient and established strains occurring at low levels (Fig. 2A). This aligns with the observed strain diversity of 1–3 co-circulating lineages (*38*). The model further suggests that influenza dynamics are dominated by rapid replacement of strains, with transient strains lasting less than 10 years (Fig.2 B,C,E), consistent with empirical observations of strain replacement within 2-5 years (*38*). The model also suggests that under conditions of elevated transmission (e.g., due to increased host density, *N*_*h*_ or higher *ε* via host birth or migration), multiple co-circulating strains can still emerge and persist (Fig. 2C).

**Figure 2.**
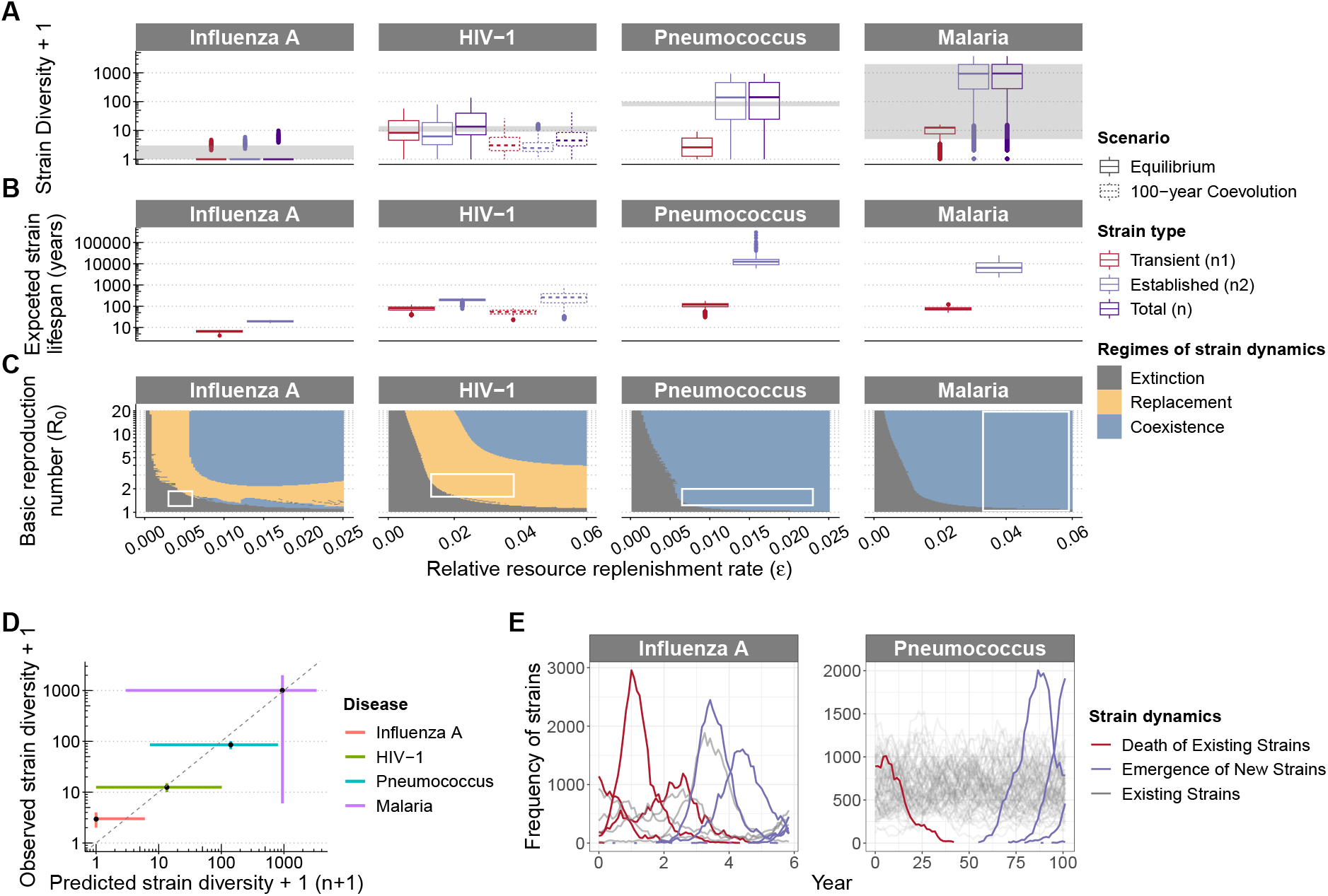
Predicted strain diversity and regimes of strain dynamics of four human diseases using MultiSEED. **(A)** Predicted numbers of transient 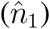, established 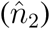, and total strains 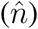 at long-term equilibrium. The explored parameter ranges of (*ε, R*_0_) combinations are shown as white rectangles in **(C)** (see Table 1 for details). Strain diversity is set to 0 when *n*_1_ + *n*_2_ *<* 1. For HIV-1, we further calculate the expected total strain diversity after 100 years of coevolution. Specifically, mean diversity was obtained from over 1,000 compartmental Markov simulations per (*ε, R*_0_) pair. Empirical strain diversity is shown as gray shades. **(B)** Corresponding strain lifespan estimates. The lifespan of a transient strain is defined as the mean duration of its initial epidemic (*T*_*e*_; see SM). The lifespan of an established strain is calculated based on its probabilities of short-term burnout and long-term persistence, using the expression 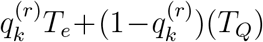 (see SM section S1.5). Cases of zero strain diversity are excluded. **(C)** Regimes of strain dynamics predicted by MultiSEED. Extinction occurs when both *n*_1_ = *n*_2_ = 0 at equilibrium. Replacement and coexistence are distinguished using two ratios: when both 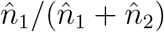 and *T*_*e*_*/*(*T*_*e*_ + *T*_*Q*_) are below 1/3, strains are considered to stably coexist; otherwise, they fall under the replacement regime. White rectangles denote empirical parameter ranges. **(D)** Comparison of MultiSEED-predicted strain diversity with observed strain diversity. For each disease, the point indicates the intersection of median predicted strain diversity across (*ε, R*_0_) combinations and the midpoint of observed range of strain diversity. The horizontal bars show the 5th-95th percentiles of predicted values, and the vertical bars show the observed range. **(E)** Representative strain dynamics for influenza A and pneumococcus using agent-based stochastic simulations, illustrating typical patterns of the replacement and coexistence regimes, respectively. See Fig. 1 legend for the parameter sets used in the simulations.

For HIV-1, the predicted equilibrium diversity is roughly an order of magnitude higher than for influenza, with the numbers of both transient and established strains ranging between 10^1^ and 10^2^. However, given HIV-1’s relatively recent emergence in humans (≈ 100 years ago) (*39, 40*), the virus is unlikely to have reached equilibrium of strain dynamics. We therefore simulated 100-year stochastic trajectories using the compartmental Markov process based on Eq. S32 (see SM Section S1.6), which yielded narrower and lower strain diversity distributions than equilibrium predictions (boxes with dashed lines in Fig. 2A). Both the diversity estimates from the equilibrium states and the 100-coevolution encompass current empirical estimates of approximately 13 known types, comprising four major groups and nine subtypes within group M (*41*) (Fig. 2D). Although HIV-1 is also in the replacement regime, its substantially higher diversity relative to influenza A and its long lifespan even for transient strains (i.e., *T*_*e*_ ≈ decades to centuries, Fig. 2B) may seem similar to the coexistence regime. Nevertheless, because most strains are still in the early phases of spread with respect to *T*_*e*_, their frequencies fluctuate more dramatically than would be expected under true long-term coexistence. These characteristics are consistent with observed phylodynamic patterns in HIV-1: high population-level diversity is maintained, while within-host evolution is continuous and often marked by reversions to ancestral strains (*42*), leading to relatively stable strain composition. Additionally, there is substantial regional variation in dominant subtypes across the globe (*43*), further supporting the picture of a dynamic yet structured strain landscape.

For pneumococcus, MultiSEED consistently predicts low numbers of transient strains (≈ 1) but a wide range of established strain diversity (from 10^1^ to 10^3^), primarily driven by small changes in *R*_0_. This prediction encompasses the roughly 90 known immunologically distinct serotypes, defined by variation in the polysaccharide capsule (*6*). Lifespan estimates show that established strains can last for millennia, placing pneumococcus firmly in the coexistence regime (Fig. 2C and E). The coexistence regime matches empirical observations: pneumococcal serotype diversity is high, stable over time and geography, and shaped by frequency-dependent selection driven by host immunity (*6*).

For malaria, we explored a broad range of *R*_0_ values associated with vector-borne transmission, which vary significantly with mosquito behavior and ecology. Consequently, predicted strain diversity spans multiple orders of magnitude, with established strains dominating the overall diversity. These predictions are consistent with the large variability in malaria parasite diversity observed across biogeographic regions with differing transmission intensity (*44*).

Although recombination continually reshuffles strain compositions, empirical evidence further supports long-term persistence of antigenic alleles (*44, 45*), genes (*46*), and modules of strains (*47*). For *var* genes, encoding the major variant surface antigens, trans-specific polymorphisms have been maintained between gorilla-associated *Plasmodium praefalciparum* and *P. falciparum*, predating *P. falciparum*’s transfer to humans ≈50,000 years ago (*44, 45*). Analyses of other major antigens also reveal high within-population antigenic diversity, high divergence among haplotypes, and little geographic structure (*46*). In addition, clusters of malaria strains have been shown to persist over extended periods, despite gradual shifts in gene composition due to recombination (*47*). Therefore, while MultiSEED does not account for sexual recombination, its predictions of strain coexistence regime are robust to this limitation.

When comparing predictions under the same parameter ranges, the AF-model (*9*) consistently overestimates strain diversity for influenza A and underestimates it for malaria (Fig. S6). Across all four diseases, the predicted diversity from the AF-model spans only about one order of magnitude, in contrast to the much broader range captured by MultiSEED. The inflation of influenza A diversity arises because the AF-model ignores extinctions during epidemic troughs, thereby overestimating the invasion success of new strains and neglecting the stochastic loss of resident strains during invasion (Fig. 1D). Conversely, by omitting the effects of mixed infections, the AF-model underestimates the hyperdiversity of strains maintained in malaria populations.

In summary, strain diversity ranges from near extinction or only a few lineages in influenza A to thousands in malaria, with observed values for all four diseases falling within the ranges predicted by MultiSEED (Fig. 2A and D). The model also reveals clear contrasts in turnover regimes: Influenza A and HIV-1 are characterized by replacement dynamics on different time scales, whereas pneumococcus and malaria exhibit long-term coexistence of multiple strains (Fig. 2C, Fig. S5B, Fig. 2E). These contrasting outcomes highlight the model’s versatility in unifying empirical patterns across pathogens with vastly different life histories.

### Factors affecting strain diversity

To further identify disease characteristics that promote or limit long-term strain diversity in general, we examine how the five model parameters interactively shape pathogen evolution (Fig. 1C). Three arise from the deterministic model: the basic reproduction number *R*_0_, the relative resource replenishment rate *ε*, and the cross-strain susceptibility *ρ*, while two additional parameters, the host population size accessible to pathogens *N*_*h*_, and the strain innovation rate *ν*, describe stochastic dynamics. Together, these parameters represent fundamental aspects of pathogen ecology and evolution, encompassing life history traits (*N*_*h*_, *R*_0_, *ε*), transmission dynamics (*N*_*h*_, *R*_0_), host immune responses (*ρ, ε*), the rate of antigenic diversification (*ν*), and host demography (*N*_*h*_, *ε*). This minimal but biologically meaningful set provides a general framework for understanding pathogen strain diversity.

Our framework shows that *ε*, rather than *R*_0_, exerts the strongest effect on disease persistence 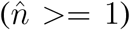, acting as a switch: below a critical threshold, no strains persist even at high *R*_0_; above it, diversity rises sharply (Fig. 3A-C, Fig S7A, Fig. S8A). Biologically, *ε* reflects the combined effects of host turnover, waning immunity, and infection duration. Thus, even modest changes in these underlying processes can profoundly alter a pathogen’s ability to sustain strain diversity over time. This threshold emerges from the inclusion of epidemic burnout in the model dynamics, making the classical condition *R*_0_ *>* 1 insufficient for sustained transmission over evolutionary timescales (Fig. 3A; see also (*22*)). The parameter region of disease extinction (lower-left grey region of Fig. 3A) represents pathogens with short infectious periods and/or long-lasting immunity, such as those causing acute infections or resulting from zoonotic spillover.

**Figure 3.**
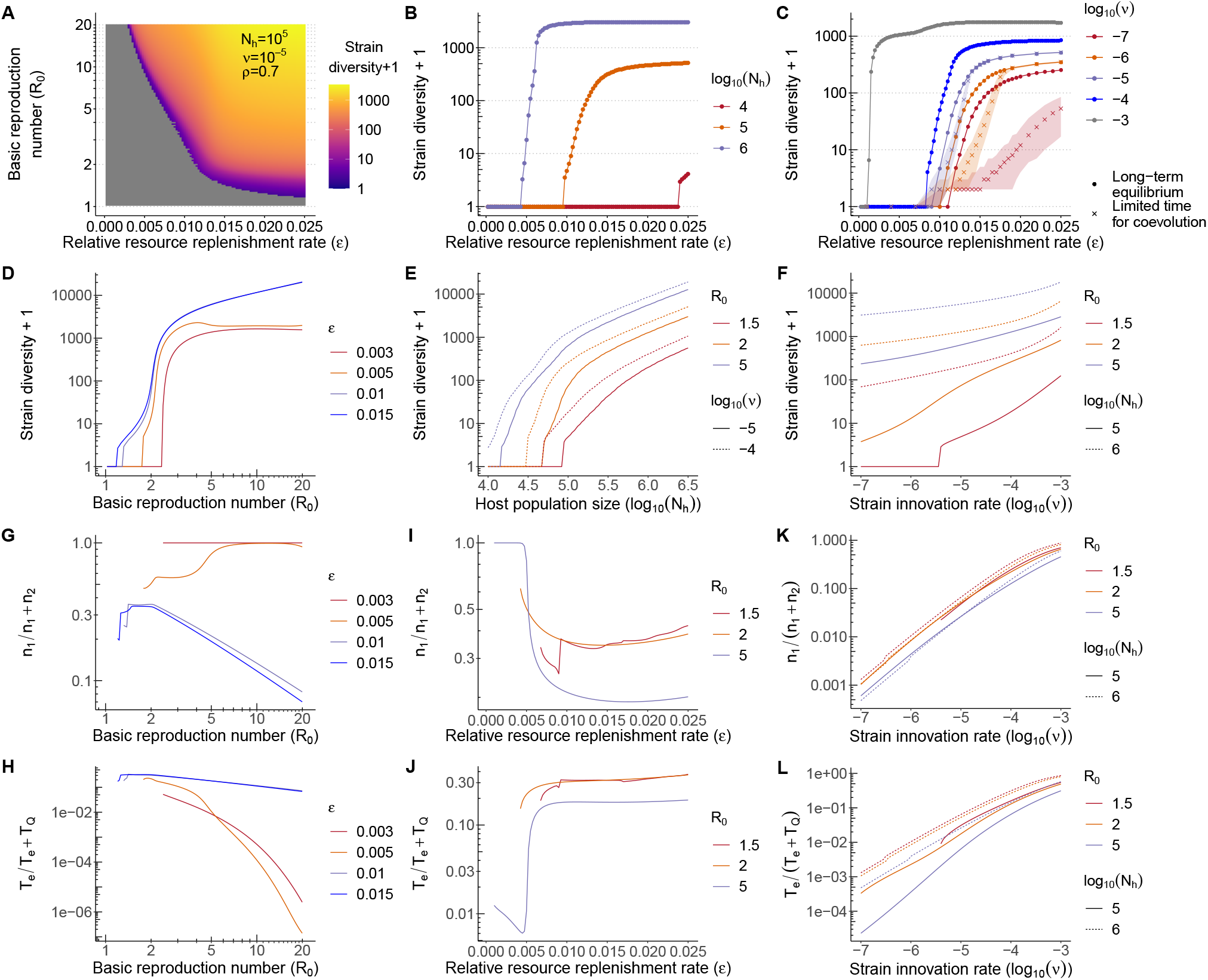
Effects of key parameters on long-term strain diversity and regimes of strain dynamics predicted by MultiSEED. **(A)** Heatmap of strain diversity showing the interplay of relative resource replenishment rate from hosts (*ε*) and the basic reproduction number (*R*_0_), with all other parameters held constant. Grid cells with zero strain diversity are shown in gray. **(B-C)** Effects of *ε* on strain diversity, across different magnitudes of host population size *N*_*h*_ and strain innovation rate *ν*. In **C**, we examine the effect of limited time for strain diversification by running stochastic compartmental model simulations for 80,000 generations, with 1,000 replicates per *ε* value (for computational efficiency, only 100 replicates were run for *ε >* 0.014 when *ν* = 10^−5^). Crosses and shades represent the median and 5th-95th percentiles across replicates. **(D)** Effect of *R*_0_ on strain diversity. Other parameters: *N*_*h*_ = 10^6^, *ν* = 10^−4^, and *ρ* = 0.5, corresponding to the parameter range for influenza A in Table 1. **(E-F)** Effects of *N*_*h*_ and *ν* on strain diversity, with fixed parameter values *ε* = 0.015 and *ρ* = 0.7. **(G-L)** Parameter effects on the two criteria used to classify regimes of strain dynamics: Effects of *R*_0_ **(G-H)** and *ε* **(I-J)**, using the same parameter settings as in **D**; Effects of *ν* **(K-L)**, with fixed parameter values *ε* = 0.015 and *ρ* = 0.7.

Once persistence is achieved, diversity grows rapidly with *R*_0_ and host population size *N*_*h*_ (Fig. 3B, D, E, Fig. S7B-D, Fig. S8B-C), but only modestly with strain innovation rate *ν* (Fig. 3C and F, Fig. S8D). Nevertheless, when the host-pathogen association is recent, increases in *ν* can significantly boost observed diversity near the persistence threshold (see shaded regions of Fig. 3C), accelerating the system’s approach to equilibrium. Thus, under conditions favorable to transmission and persistence, strain diversity is less dependent on the innovation rate, which contrasts with classical neutral models in population genetics, where *N*_*h*_ and *ν* contribute equally to diversity. In disease dynamics, large *N*_*h*_ not only increases the supply of new strains but also reduces burnout risk and extends the average lifespan (*T*_*Q*_) of established strains after successful invasion. In contrast, *ν* affects only the rate of strain emergence, not their survival. Our findings suggest that long-term diversity is governed mainly by ecological factors, including pathogen life history and host-pathogen interactions, rather than intrinsic pathogen genetic factors *ν* and cross-reactivity between antigens *ρ* (Fig. S7A-B, Fig. S8E).

### Factors influencing regimes of strain dynamics

Because strain turnover regimes depend on parameters in complex, often nonlinear ways, the factors that shape diversity are not always the same as those that govern regime classification. Notably, the two parameters that define the strain persistence threshold, *ε* and *R*_0_, do not always have monotonic effects on the strain regimes (Fig. 3G–J; see also Fig. S8A–B for additional parameter combinations). In general, increasing *ε* or *R*_0_ beyond the persistence threshold promotes long-term strain coexistence. However, when either parameter is small or persistence is close to the threshold, the system tends to favor strain replacement.

Among all parameters, the strain innovation rate *ν* has a particularly strong and consistent effect on the regimes of strain dynamics. Increases in *ν* drive a near-linear shift toward strain replacement, regardless of other parameter values (Fig. 3K–L). In contrast, host population size *N*_*h*_, while critical for determining the persistence threshold and setting the upper bound of strain diversity (Fig. 3E), has only a modest effect on regime classification. Larger *N*_*h*_ promotes strain replacement, probably due to its dual role as both a resource base for sustaining existing strains and as a reservoir for seeding new invasions (Fig. S8C). Lower cross immunity (i.e., higher *ρ*) tends to promote strain coexistence, but its effect becomes more variable near the persistence threshold (Fig. S8E), where parameter interactions become highly nonlinear. Together, these results underscore that the regimes of strain dynamics are shaped by non-additive, sometimes non-monotonic interactions among parameters, especially near the persistence threshold. As such, regime classification under specific parameter combinations must be determined numerically through MultiSEED simulations, rather than from intuition or theoretical expectations.

### Summary and Perspectives

In summary, we present a novel numerical framework for predicting long-term strain diversity and regimes of strain dynamics. By distinguishing transient from established strains, Multi-SEED reveals whether pathogens are governed by replacement or coexistence, providing insights into the qualitative nature of strain dynamics that earlier models could not capture. By considering finite strain-specific immunity and mixed infections, our framework reproduces the observed patterns of strain diversity, persistence, and turnover regimes in focal human diseases, which exhibit a wide range of life history traits and transmission modes.

One key conceptual contribution of this framework is the introduction of a composite parameter, *ε*, which represents the relative rate of resource replenishment for susceptible hosts. This parameter accounts for both immunity loss and demographic turnover relative to infection recovery rate, and emerges as a key determinant of disease persistence. Previous studies have shown that in the absence of sufficient host replenishment (e.g., susceptible individuals are replenished solely through new births in the population), long-term persistence is unlikely for most human diseases, particularly for short-duration infections (*22*). Our results expand on this by demonstrating that finite strain-specific immunity significantly boosts *ε* and ensures persistence or promotes high strain diversity, as seen in influenza A and malaria, respectively (compare white rectangles with black bars in Fig. S4A). The persistence filter effect of *ε* also impacts critical community sizes (CCS), a classical measure of disease persistence. Previous studies derived CCS from the estimated long-term lifespans of strains (i.e., *T*_*Q*_ in our model) (*9, 36, 48*). Our study suggests that the required CCS would be higher, because the pathogen needs to survive the burnout stage in addition to enduring long-term stochastic loss.

Our framework confirms that increases in any of the five key parameters lead to greater long-term strain diversity (*9, 10, 18, 49*), while also quantifying the relative importance of each parameter in shaping that diversity. In general, we show that host population size (*N*_*h*_) and *R*_0_ have greater influences on equilibrium diversity. When close to the persistence threshold with small *R*_0_, parameters such as cross-immunity (1-*ρ*) and innovation rate (*ν*) become more critical, where a small reduction in cross-immunity can lead to large shifts in diversity magnitudes.

Unlike strain diversity, the impacts of parameters on strain dynamics are often non-monotonic, emerging from complex interactions among multiple parameters, especially near the persistence threshold. This complexity is exemplified by comparing the dynamics of influenza A and pneumococcus. Despite similar values of *R*_0_, *N*_*h*_, and overlapping ranges of *ε*, pneumococcus exhibits stable coexistence, while influenza A displays frequent strain replacement. This divergence is largely driven by lower cross-immunity and lower innovation rate of pneumococcus, which permit long-term persistence of strains (Table 1). Our results also challenge earlier explanations that invoked low mutation rates (*12, 13*) or transient cross-immunity (*14*) to explain influenza A’s low diversity and replacement regime. Instead, MultiSEED identifies the combination of low *R*_0_ and low *ε* as the main constraint on influenza diversity, while its higher mutation rate, relative to bacterial or protist pathogens, drives continual strain replacement.

Our framework also provides a powerful tool for studying how pathogen diversity may respond to environmental and demographic changes. Global shifts such as increasing human mobility, urbanization, or climate-induced habitat change can affect key epidemiological parameters like *R*_0_, *N*_*h*_, and *ε*, with potentially large effects on strain diversity and turnover. For example, greater connectivity among human populations may increase effective transmissibility and host replenishment, leading to larger *R*_0_ and *ε*, which in turn increase strain diversity. Meanwhile, public health interventions, such as vaccination campaigns, will reduce the susceptible pool and the immune loss rate, thereby decreasing the host population size *N*_*h*_ and the resource replenishment rate *ε*, lowering the likelihood of diverse strain coexistence. MultiSEED can be applied to evaluate how such global trends influence pathogen evolution and to inform strategies for disease control in an increasingly interconnected world.

While MultiSEED captures a wide range of multi-strain dynamics, it also makes several simplifying assumptions. First, it assumes that all strains are epidemiologically equivalent, with equal transmissibility, recovery rates, and immune waning. In reality, fitness differences among strains can also affect their invasion probabilities and persistence (*19*). However, recent simulation work suggests that under strong immune-mediated competition, functional fitness across strains tends to converge over evolutionary timescales while immunological niche differences are widened (*50*). We thus predict that the equal fitness assumption is more valid in high-diversity systems. In contrast, in transmission-limited or low-diversity scenarios, deviations from this assumption may lead MultiSEED to overestimate the number of coexisting strains. A natural extension would be to allow for two or more classes of strains with differing functional fitness. Second, the current framework is non-spatial and does not incorporate host contact structures. In reality, pathogens that fall into the extinction regime could still persist over evolutionary timescales through asynchronous epidemics in spatially structured populations, as observed for measles (*51*). The value of the non-spatial approach lies in highlighting the underlying mechanism of low diversity in measles: when *ε* is very low (i.e., a short disease duration and no immune memory loss), even a high *R*_0_ leads to long-term local extinction and thus low diversity. By contrast, the AF-model would predict extremely high diversity in thousands under the same conditions of low *ε* and high *R*_0_ (Fig. S6A). Third, the current model assumes a single host species, whereas many pathogens infect multiple hosts. A larger host range will increase *ε* and *N*_*h*_, promoting established strain diversity in density-dependent transmission, or buffer transmission due to a reduction in *R*_0_ (*52*). Future work could incorporate multi-host systems with varying susceptibilities and transmission roles. Lastly, the SIRS framework may not apply to all pathogens. For instance, tuberculosis involves complex latency dynamics (*53*), while dengue virus features immune enhancement upon reinfection (*54*); however, these interactions are not captured in the current model. Thus, the deterministic model in MultiSEED would likely have to be adapted for these specific transmission systems before its application. Additionally, we have explored parameter values relevant to human diseases. Applying MultiSEED to plant, animal, or vector-borne pathogens would require adapting assumptions about host demography, immune architecture, and transmission biology to reflect system-specific constraints and mechanisms.

In conclusion, MultiSEED represents a significant step forward in understanding the ecoevolutionary drivers of pathogen diversity and strain dynamics. The model not only reproduces empirical observations in multiple human diseases, but also provides a flexible framework for evaluating how pathogen diversity may respond to global change, demographic shifts, and intervention strategies. Future developments include accommodating fitness variation of strains, multi-host dynamics, and investigating different types of transmission dynamics.

## Acknowledgments

We thank Greg Dwyer and Mercedes Pascual for their help in improving the manuscript. We are thankful to Matthias Steinruecken and Samy Tindel for insightful discussions regarding model construction. We thank Pradyut Kumar for his help with the figure illustrations. We appreciate the High-Performance Computing support from the Rosen Center for Advanced Computing at Purdue University and the Purdue Institute of Inflammation, Immunology, and Infectious Disease.

## Authors contributions

J.L., methodology, data curation, software, formal analysis, visualization and writing; Q.H., conceptualization, methodology, software, formal analysis, visualization and writing.

## Data and materials availability

All data and code are available at GitHub: https://github.itap.purdue.edu/HeLab/MultiSEED

## Supplementary Materials

Materials and Methods

Supplementary Text

Figs. S1 to S10

Table S1

References (108-118)

## Supplementary Materials

## 1 Material and Methods

In the following sections, we introduce the construction of the MultiSEED framework (Fig. S1). We start by extending a single-strain SIRS model to a status-based *n*-strain SIRS model (Fig. S1A, Section 1.1-1.2), followed by a detailed description of the system’s stochastic dynamics at ecological and evolutionary time scales (Section 1.3-1.4). We then introduce the numerical methods for obtaining long-term strain diversity by defining the MultiSEED compartmental model of strain dynamics (Fig. S1B, Section 1.5-1.6). Results from the numerical solutions are verified with simulations using an agent-based stochastic model (Section 1.7). Parameter ranges of interest are explored using the empirical ranges obtained from common infectious diseases (Section 1.8).

### 1.1 Single-strain SIRS dynamics

For the SIRS model of a disease with vital dynamics, we use *S, I, R* to denote the number of susceptible, infected, and recovered people in the total population. Birth and death rates of the hosts are equal, such that the total host population size, *N*_*h*_ = *S*(*t*) + *I*(*t*) + *R*(*t*), is assumed to be constant. Then the deterministic mean-field ODEs are,

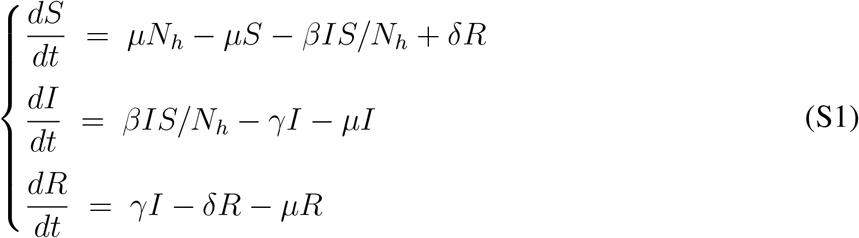

where *µ* is the *per capita* natural birth or death rate of the host population, *β* is the transmission rate, *γ* is the recovery rate from the infection, and *δ* is the immunity loss rate of the recovered class.

If we use *X, Y*, and *Z* to denote the fractions of susceptible, infected and recovered (i.e., 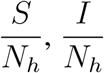, and 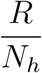), we can rewrite Eq. S1 to be,

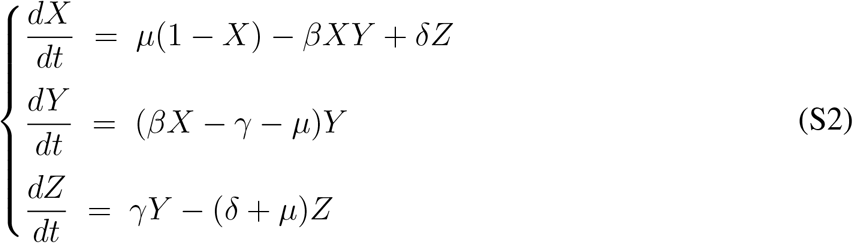

Given that 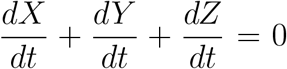 and *X* + *Y* + *Z* = 1 being constant, we may study the first two equations to describe the system,

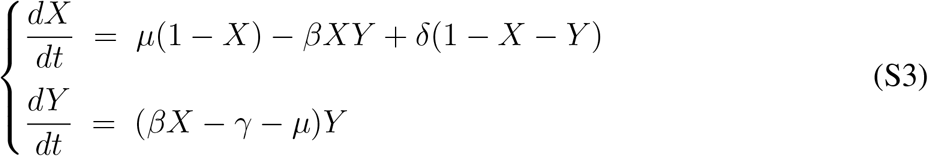

The trajectories of the deterministic SIRS model converge to a globally asymptotically stable equilibrium point, which is determined by the basic reproduction number, 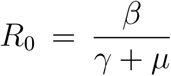. When *R*_0_ *<*= 1, the system will be disease-free, whereas *R*_0_ *>* 1, the system goes to an endemic equilibrium,

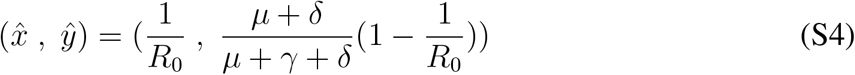

If we reparameterize the system and rescale time using the expected duration of an individual infection, we define a new time variable *τ* = *t/*(*γ* + *µ*), Eq S3 becomes,

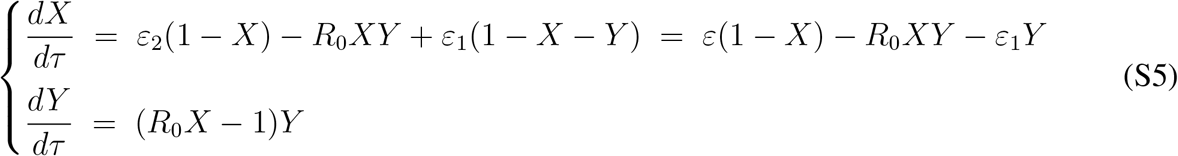

where 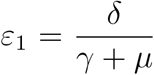 represents the rescaled immune loss rate in the units of expected infectious period 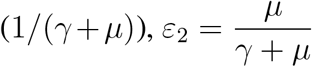 represents the rescaled birth and death rates of hosts in the units of expected infectious period, and *ε* = *ε*_1_ + *ε*_2_.

### 1.2 Multi-strain SIRS dynamics

#### 1.2.1 status-based *n*-strain SIRS model

Many infectious diseases consist of multiple co-circulating strains, and hosts may be infected by different strains either simultaneously or sequentially. In this study, we extend the classic single-strain SIRS framework to model the dynamics of multiple strains. We assume there are currently *n* equivalent strains in circulation, denoted as *H* = {1, 2, …, *n*}, where all strains share identical epidemiological properties, including *R*_0_ and immune loss rate (*δ*). For simplicity, we initially do not incorporate cross-immunity between strains, but we extend the model in the next section.

Unlike earlier multi-strain models that restrict hosts to infection by only one strain at a time (such as (*9*)), we allow for simultaneous co-infection by multiple strains, reflecting the common occurrence of mixed infections observed in empirical studies (*27*). In our model, strains compete within co-infected hosts by sharing transmission potential: if *k* different strains co-infect the same host, the transmissibility of each strain is reduced to *β/k*. We also assume that there is no trade-off between transmissibility and virulence, so the recovery rate *γ* of each strain within the host remains unchanged from that of single-strain infections.

We do not track the full infection history of individual hosts. Instead, we record their current immunity and infection status. This allows us to continue using Eq. S5 to describe the transmission dynamics of individual strains, with the exception that the transmission rate *β* and the corresponding 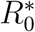 of strains must be adjusted to account for co-infection, where multiple strains share the same host. Let *x*_*i*_(*τ*), *y*_*i*_(*τ*), and *z*_*i*_(*τ*) denote the proportions of hosts at time *τ* who are susceptible to, infected by, and recovered from strain *i*, respectively. By definition, these proportions satisfy *x*_*i*_(*τ*)+*y*_*i*_(*τ*)+*z*_*i*_(*τ*) = 1 for each strain *i*. However, since hosts can be co-infected with multiple strains, the total infected proportion across all strains, 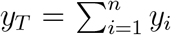 can exceed 1. Assuming no host age structure and that co-infections are randomly distributed among hosts, we define the overall disease prevalence (i.e., the probability that a host is infected with at least one strain) as 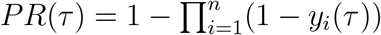.

Assume that 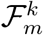 represents the *k*th random subset of *m* strains out of *H\i*, the total number of such subsets is *C*(*n* − 1, *m*). Then, the reduced transmissibility of strain *i* because of coinfection with other strains can be calculated as follows,

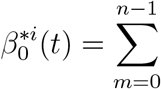 (fraction of *β*|(*m* + 1) infections) · *P* (Host having *m* infections in addition to strain *i*)

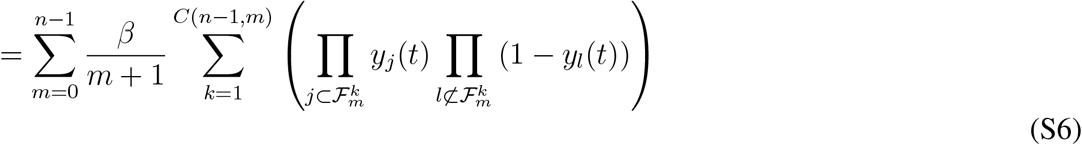

Therefore, we can modify Eq. S5 to describe the dynamics of a particular strain within *n* equivalent circulating strains to be,

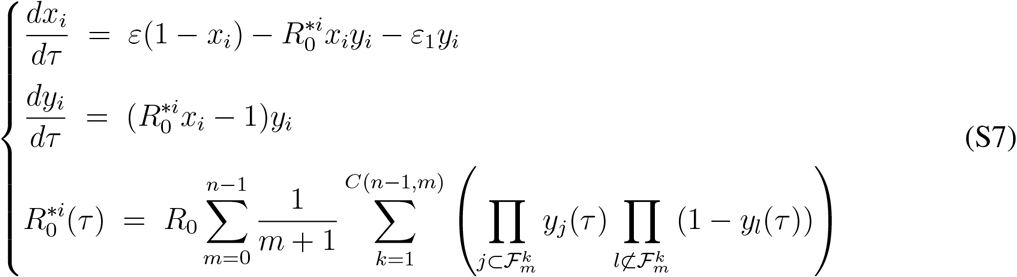

In quasi-stationary state, each strain is roughly of equal frequency so that *ŷ*_*i*_ = *ŷ*_*T*_ */n*. We can then approximate Eq. S6 as follows,

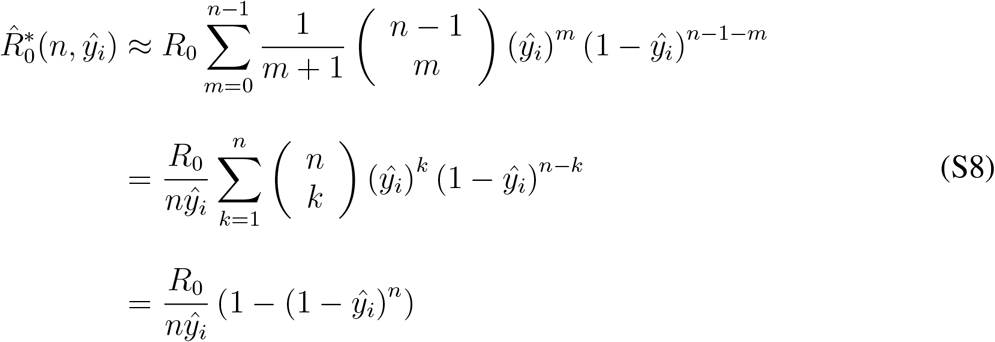

We can thus solve for the equilibrium values 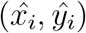 and 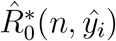 at the quasi-stationary states.

#### 1.2.2 Incorporating generalized cross-immunity

In the previous section, we assumed that there was no cross-immunity among strains. However, in reality, prior infections often confer partial protection against subsequent heterologous infections. To capture this, we now consider an extended version of *n*-strain models that incorporates general cross-immunity. We adopt the definition of cross-immunity from (*108*) and generalize it to accommodate multiple strains. For tractability, we assume that hosts with any prior infection history gain a constant level of cross-protection to all strains in the form of reduced susceptibility. Hosts do not lose generalized immunity. While hosts may lose strain-specific immunity and re-enter the susceptible class for that particular strain, they still maintain the generalized protection, modeled as a reduced susceptibility. Accordingly, we modify Eq. S7 for *n* equivalent strains as follows:

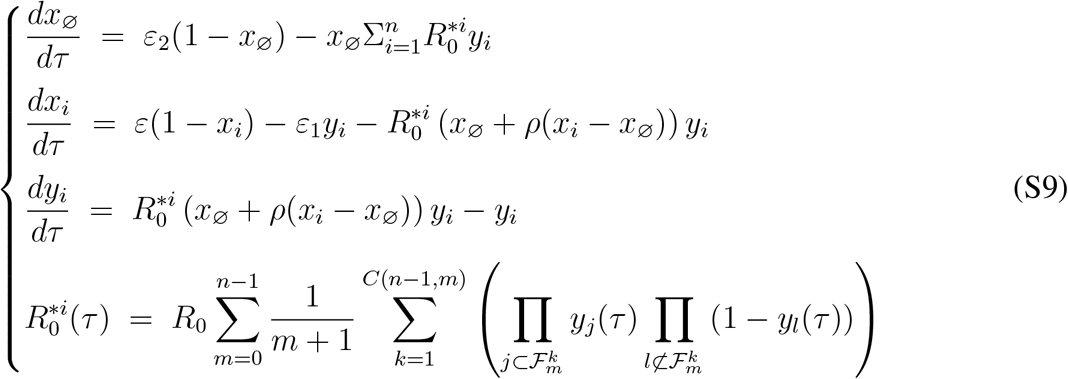

Here, the definitions of *x*_*i*_, *y*_*i*_, and 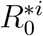 remain consistent with those in Eq. S7. We specifically track the dynamics of *x*_∅_, the proportion of naïve hosts who have never been infected by any strain and are therefore fully susceptible to all strains, without any cross-protection. *x*_∅_ will increase only through host birth, and will decrease from host deaths and infection by any strain. The parameter *ρ* quantifies cross-susceptibility of non-naïve hosts who are still susceptible to strain *i, x*_*i\*∅_. These hosts include the ones infected by strains other than *i*, or those who have lost specific immunity to strain *i*. Thus, *x*_*i\*∅_ = *x*_*i*_ − *x*_∅_. A value of *ρ* = 1 corresponds to no cross-immunity, whereas *ρ* = 0 indicates complete cross-protection. We again use Eq. S8 and Eq. S9 to solve for the equilibrium values 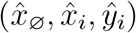 at the quasi-stationary state.

#### 1.2.3 Deterministic dynamics of a new-strain invasion into *n* existing strains at equilibrium

We have established the equilibrium dynamics of *n* strains. To explore the invasion potential by a novel strain, we extend our ODEs to include the dynamics of a single invading strain, *ω*, introduced into a parasite population where the *n* resident strains are at equilibrium. We first present the deterministic ODEs governing this invasion process. In the next sections, we incorporate stochastic dynamics to estimate the probability of successful invasion.

Let *x*_*r*_ and *y*_*r*_ represent the susceptible and infected compartments associated with each of the *n* resident strains, respectively, and *x*_*ω*_ and *y*_*ω*_ denote the corresponding compartments for the invading strain *ω*. The following system of ODEs describes the deterministic dynamics of this invasion scenario.

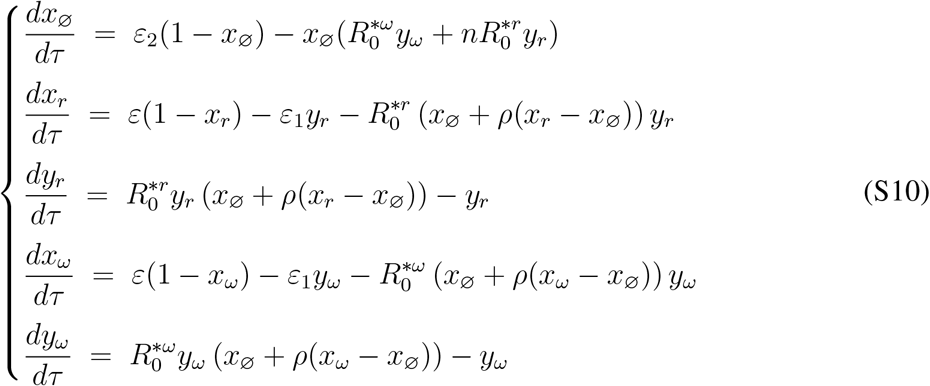

Here *x*_∅_ is still the fraction of susceptible hosts with no previous infection, while 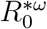 and 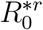 refer to the effective reproductive number of the invading strain and the resident strains, respectively. Similar to Eq. S8, the reduced transmissibility of each resident strain is due to coinfection with other resident strains and/or the new invading strain. Thus,

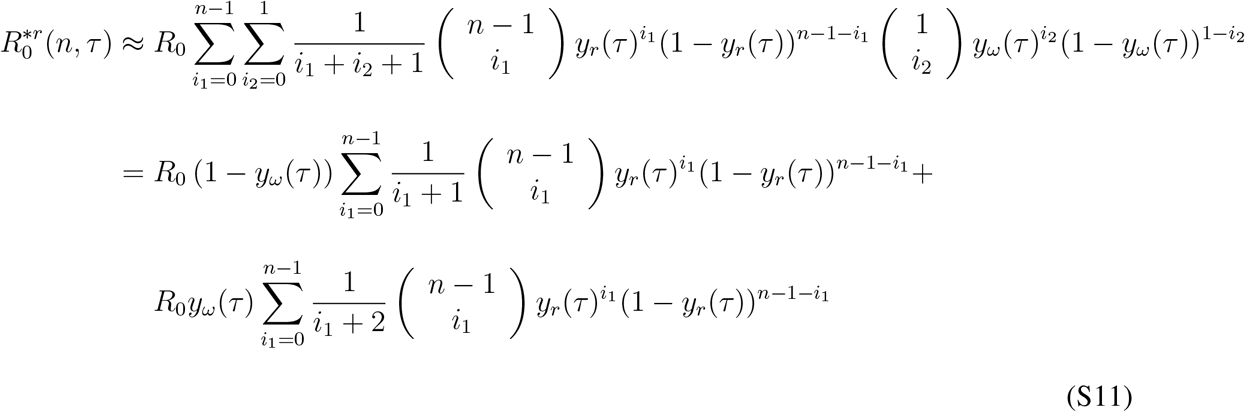

The second term of Eq. S11 is equivalent to solving 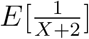 given *X* is a random variable following a binomial distribution *B*(*n* − 1, *y*_*r*_(*τ*)). Using generating functions, we get a closed form solution for 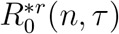,

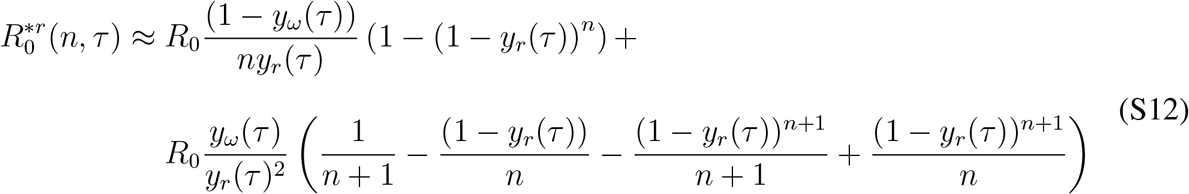

The reduced transmissibility of the new invading strain, 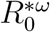, on the other hand, is due to the competition with the *n* resident strains. It can be derived in a similar way to Eq. S8:

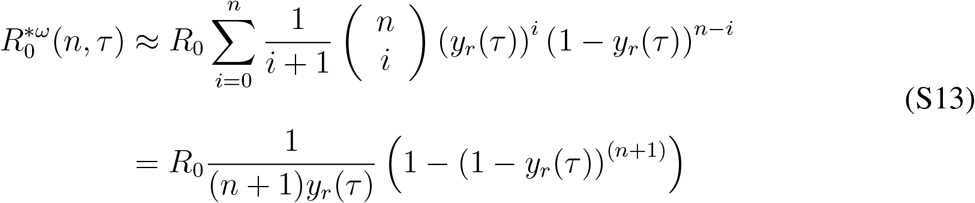

### 1.3 Continuous time stochastic processes at the ecological time scale: rates of strain emergence and strain dynamics

To predict the number of strains at the long-term equilibrium, the next step is to quantify the dynamics of strain emergence and extinction. A newly formed strain has to survive two phases of bottlenecks to become a established strain. Strains are mostly prone to extinction during the initial fizzle phase, when a small number of infections may stochastically go extinct. Once the strain accumulates enough copies and reaches the epidemic stage, it does not go extinct until the susceptibles are quickly exhausted and the fraction of infected individuals again decreases sharply, a phenomenon termed epidemic burnout (see trends of light red lines between *τ*_1_ and *τ*_2_ in Fig. S2). Meanwhile, a new epidemic also affects the dynamics of *n* resident strains, as the fractions of resident strains, *y*_*r*_, decrease, resulting in resident strain burnout (see trends of light blue lines between *τ*_3_ and *τ*_4_ in Fig. S2).

We build on the boundary layer framework for SIR models developed in (*21*) and (*22*) to estimate short-term extinction probabilities—specifically, the fizzle and burnout rates—following the introduction of a new strain in an *n*-strain SIRS model. Using the deterministic ODE trajectories described in Section 1.2.3, we approximate the epidemic phase until the prevalence of the invading strain *ω* and any resident strain *r*, denoted *y*_*ω*_ and *y*_*r*_, fall below a threshold *ŷ*_*n*+1_. This threshold defines a boundary layer below which stochastic effects dominate. Within this stochastic regime, we apply Kendall’s *q* function (*109*), which models a time-inhomogeneous birth-death process, to estimate the extinction probabilities for strains whose prevalence has dropped below *ŷ*_*n*+1_.

Following the initial epidemic wave, surviving strains may undergo additional cycles of resurgence and decline near *ŷ*_*n*+1_. However, (*22*) showed that the probability of extinction after surviving the first trough is negligible compared to extinction during the initial wave. Therefore, we restrict our analysis to the extinction probabilities resulting from the first epidemic wave to approximate the short-term fizzle and burnout rates for both invading and resident strains.

According to Kendall (*109*), in a time-inhomogeneous birth-death process with per capita birth rate of *b*(*τ*) and death rate of *d*(*τ*), the probability of the dynamics starting from one individual is extinct by the time *τ, q*(*τ*) is,

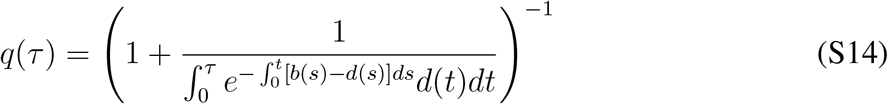

#### Fizzle stage

Assume that currently the system has *n* strains at the endemic stage. Recall the dynamics of *y*_*ω*_ is,

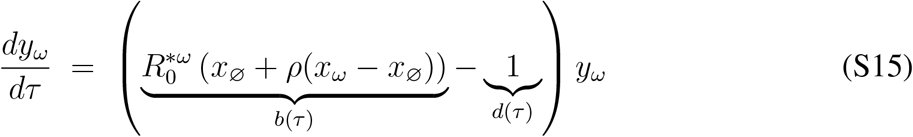

The per capita birth rate of *ω* is 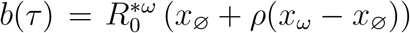, and the per capita death rate is *d*(*τ*) = 1. At the beginning of strain emergence (i.e., when *τ* is small), 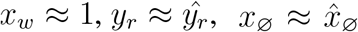 at *n*-strain equilibrium. Thus 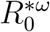 at *τ* = 0 can be considered constant at the fizzle stage as well. Therefore, the probability of the new strain *ω* escaping the fizzle stage is,

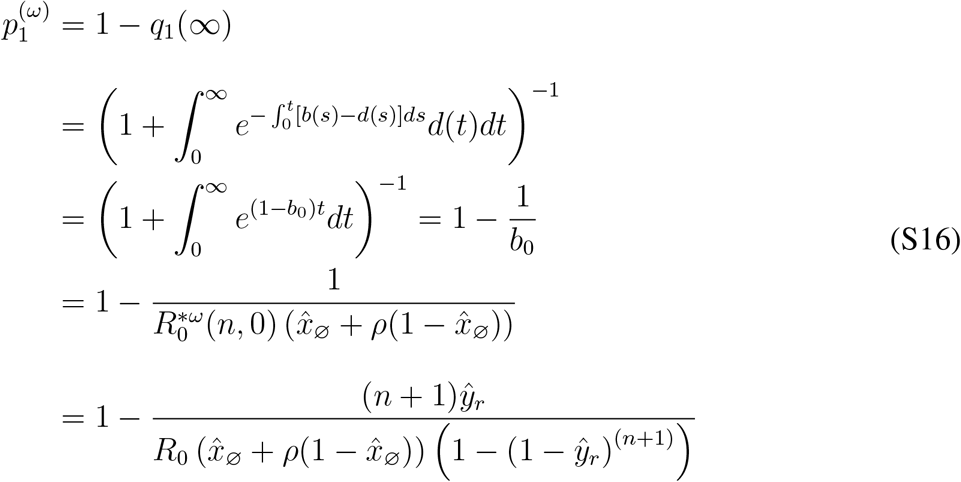

For the standard single strain SIR model, *b*_0_ = *R*_0_, and 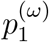 is equivalent to the classical expression for the establishment probability of a new disease 1 − (1*/R*_0_)^*k*^, where *k* is *I*(0), the initial infectious individuals (*35*).

#### Burnout phase

Once a newly formed strain survives the fizzle phase, it enters its epidemic stage and reaches a peak of infection. We use the numerical solutions of the deterministic ODEs in Eq. S10 to approximate the trajectory of strain *ω* (Fig. S2). When the fraction of infected declines and re-enters the boundary layer (i.e., drops below the threshold *ŷ*_*n*+1_), we again use Kendall’s *q* (Eq. S14) to estimate burnout probability between *τ*_1_ and *τ*_2_ for the invading strain, *ω*, and between *τ*_3_ and *τ*_4_ for the resident strains *r* (Fig. S2).

Within the boundary layer, the per capita death rate remains constant at 1, while the per capita birth rate, *b*(*τ*) varies dynamically due to rapid changes in 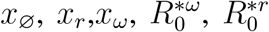. We numerically solve the ODEs to obtain the deterministic time series for these variables and calculate the corresponding *b*(*τ*) using Eq. S12 and Eq. S13, which serve as inputs to Kendall’s *q* function. The critical times *τ*_1_, *τ*_2_,*τ*_3_ and *τ*_4_ are determined by when the prevalence trajectories intersect with the boundary threshold *ŷ*_*n*+1_ (Fig. S2). We then numerically integrate Kendall’s *q* to approximate the per infection burnout probability for the invading and resident strains respectively. Upon entering the boundary layer, the strains have *k* = *N*_*h*_*ŷ*_*n*+1_ individuals. Thus the final burnout probability is given by 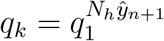.

n probability of the infection starting from *k* = *N*_*h*_*ŷ* individuals. At time *τ*, as shown in Eq. S5, the birth rate of the infected class *y* is *b*(*τ*) = *R*_0_*X*(*τ*), and the death rate of the infected class is *d*(*τ*) = 1. In the multi-strain scenario, for each strain *i* we need to find out the real-time adjusted 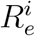 and fraction of susceptible hosts *x*_*i*_ when the fraction of infected hosts *y*_*i*_ is below *ŷ*, and then we can calculate the extinction probability using Eq. S14.

In the special case where no resident strains are present, the invading strain is the sole circulating strain. The ODE system simplifies to Eq. S17:

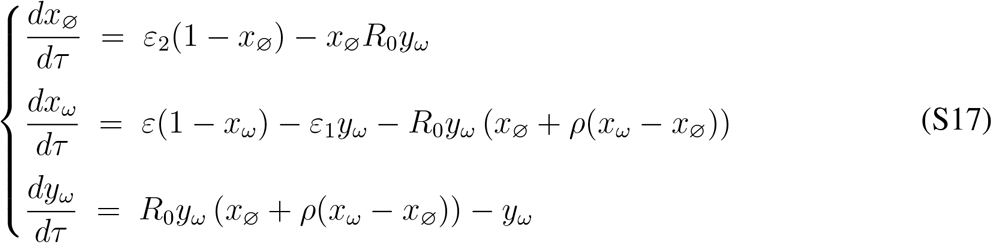

We follow the same steps as mentioned above to calculate the fizzle and burnout probabilities of this new strain after its first epidemic. In this scenario, there is no risk of burnout for the non-existent established strains.

Another key quantity we aim to estimate is the mean duration of the initial epidemic, *T*_*e*_, for invading strains that go extinct during the burnout phase. This metric helps quantify the sojourn time of newly introduced strains. As explained earlier, we use the deterministic processes to approximate the epidemic phase from *τ* = 0 to *τ*_1_ (Fig. S2). Between *τ*_1_ to *τ*_2_, we apply Kendall’s *q*-function to estimate the mean time to extinction, *τ*_*e*_. To further characterize strain dynamics between *τ*_1_ and *τ*_2_, we first calculate the mean duration of strain presence—accounting for both extinct and surviving strains—using the survival probability function *q*(*τ*). Let *q*_*k*_(*τ*) represent the probability that a strain with *k* initial copies has gone extinct by time *τ*. Then, the expected mean duration of presence across this interval is:

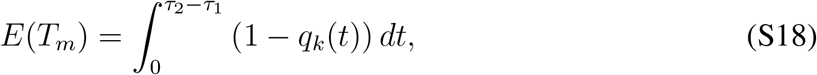

where 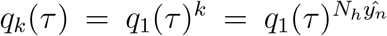. Note that this integral does not have a closed-form solution but can be approximated numerically using ODE solvers. For strains that survive the period, the mean duration is simply *τ*_2_ − *τ*_1_. Combining the contributions from both extinct and surviving strains, we obtain *E*(*T*_*m*_) = *τ*_*e*_ · *q*_*k*_(*τ*_2_ − *τ*_1_) + (*τ*_2_ − *τ*_1_) · (1 − *q*_*k*_(*τ*_2_ − *τ*_1_)). Solving for *τ*_*e*_, we rearrange the expression:

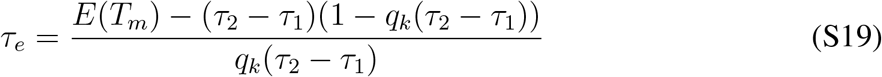

Thus, the total expected duration of the first epidemic is given by,

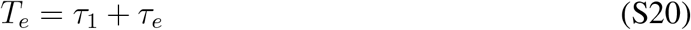

### 1.4 Continuous time stochastic processes at the evolutionary time scale: time to extinction of an endemic strain

If an infectious disease strain survives the initial troughs and stays around the endemic equilibrium 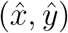, then how long does it take for it to go extinct stochastically? Due to negative frequency-dependent selection, the prevalence of the strain will be maintained at the endemic level, appearing to be in a stable equilibrium before eventually becoming extinct. The sojourn time depends on *ŷ* as well as the intensity of competition against other co-circulating strains. We write out the Kolmogorov forward (aka Fokker-Planck) equation for the *n*-strain SIRS model and derive the joint quasi-stationary distributions of *I*_*i*_ and *S*_*i*_ at time *τ, p*_*mn*_(*τ*) = *P* (*S*_*i*_(*τ*) = *m, I*_*i*_(*τ*) = *n*). We then use a bivariate Ornstein-Uhlenbeck process to approximate the distribution. The expected sojourn time to extinction is thus the inverse of the probability of having one infected host in the quasi-stationary state times the duration of an individual infection (Eq. 2.8 in (*36*)).

First, we write out all the transition rates for the stochastic version of the one-strain SIRS model. We can then calculate mean, variance, and covariance of *x*(*τ*) and *y*(*τ*) considering a bivariate diffusion approximation. The approach of approximating the mean sojourn time is then extended to the *n*-strain scenario.

The changes in the state variables *x* and *y* during the time interval from *τ* to *τ* + Δ*τ* are denoted Δ*x* and Δ*y*. We derive the mean and the covariance of Δ*x* and Δ*y*. The mean is,

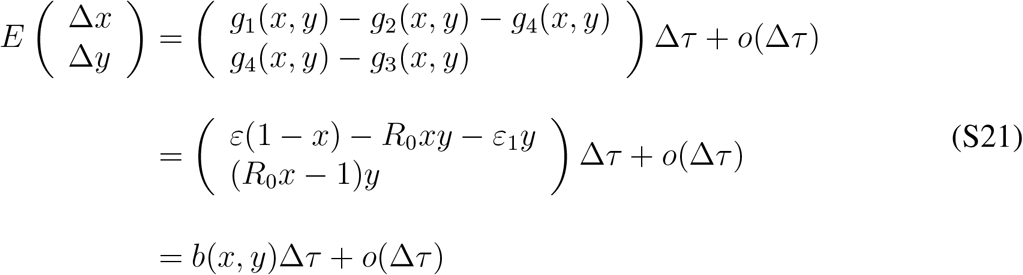

The Jacobian matrix of the vector *b*(*x, y*) with respect to *x* and *y* is denoted *B*(*x, y*):

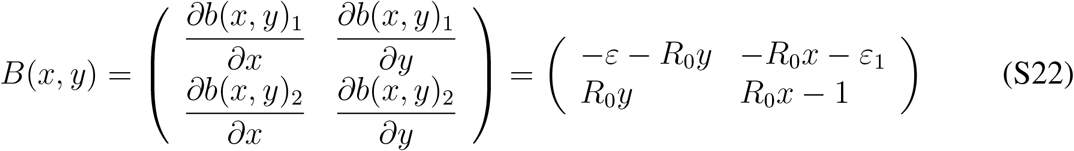

When evaluated at the endemic equilibrium point 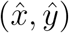, we get,

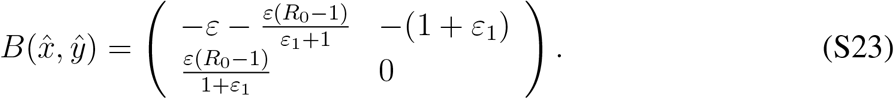

We then determine the covariance matrix of the vector of changes in the state variables during the time interval (*τ, τ* + Δ*τ*),

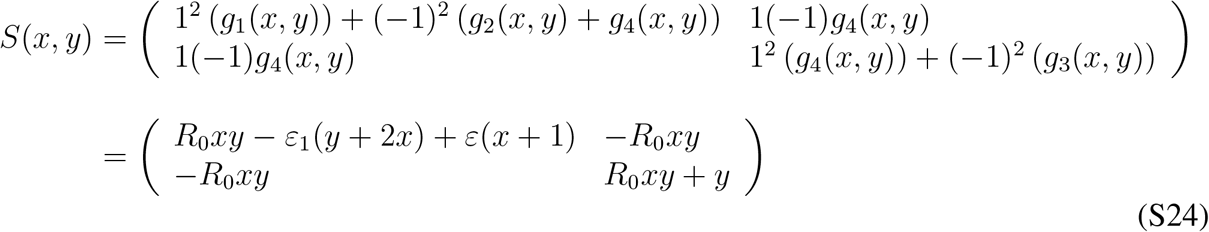

When evaluated at the endemic equilibrium point 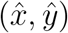, we get,

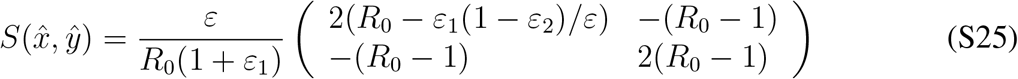

For large *N*_*h*_, the process 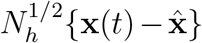 is approximated by a bivariate Orstein-Uhlenback process with local drift matrix 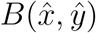 and local covariance matrix 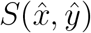. Its stationary distribution approximates the quasi-stationary distribution, which is approximately a bivariate normal distribution with mean 0 and covariance matrix Σ, where *B*Σ + Σ*B*^*T*^ = −*S* (*37*). Σ is thus,

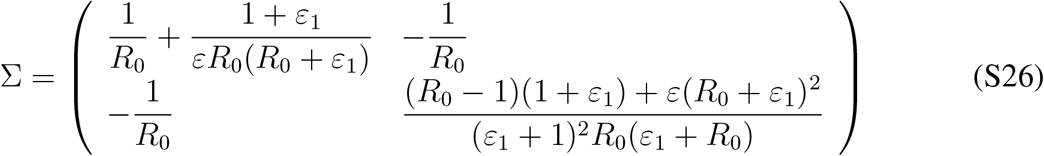

Therefore, we can approximate the marginal distribution of infected individuals *I* with the normal distribution, 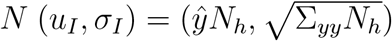. Therefore,

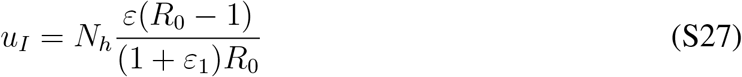

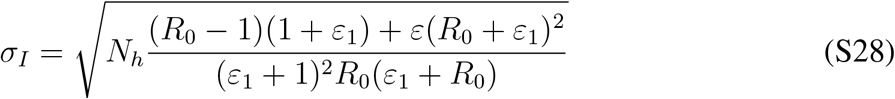

When *ε* is small, that is, the immune loss and birth rates are much smaller than the recovery rate of the disease,

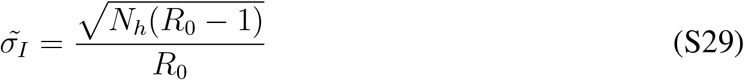

We use *f*_·1_ to denote the conditional probability of *I* = 1 given *I* being positive. That is, 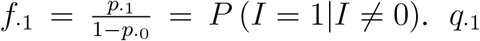 can thus be approximated using the probability density function of normal distribution rescaled by truncating the cumulative density at 0.5,

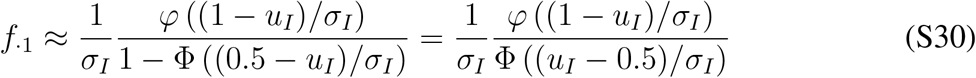

Here *φ* and φ are the PDF and CDF of the standard normal distribution. Then using Eq. 2.8 in (*36*), we get the approximation for the mean time to long-term stochastic extinction in the unit of the duration of infection,

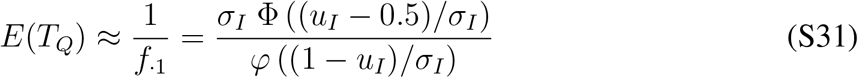

We extend the derivation of *T*_*Q*_ of one strain SIRS model to the *n*-strain model by modifying Eq. S27 and Eq. S28. Specifically, *R*_0_ in these two equations needs to be replaced by 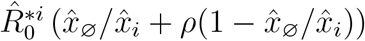 to reflect the competition among strains and the reduced susceptibility from generalized immunity.

### 1.5 Long-term strain diversity at equilibrium

In previous sections, we described the ecological and evolutionary dynamics of strains, from their emergence to extinction. Strains accumulate through the continual invasion of new mutants and are lost either during early burnout phases or through long-term stochastic extinction. To quantify strain diversity at equilibrium, we model two categories of strains: transient strains (*n*_1_) that recently emerged and have survived the initial fizzle phase but have not yet completed their first epidemic wave, and established strains (*n*_2_) that have survived at least one epidemic wave and whose long-term persistence is governed by a stochastic extinction timescale *T*_*Q*_. Consequently, total strain diversity is *n* = *n*_1_ + *n*_2_.

#### Birth and Death Dynamics of Transient Strains

Let *N*_*h*_ be the host population size and *n* = *n*_1_ + *n*_2_ be the total number of strains at time *τ*. Then the overall prevalence is given by 1 − (1 − *ŷ*)^*n*^. Assuming a strain innovation rate *ν*, the birth rate of transient strains is 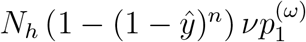, which represents the rate at which new strains are introduced and survive the initial fizzle phase. For the existing transient strains in the system, we will quantify their durations of the first epidemic to calculate the exit rate of the transient class, 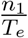, where *T*_*e*_ is the mean duration of the first epidemic, as calculated by Eq. S20.

#### Transition from transient to established strains and Extinction of established Strains

Transient strains become established at the rate of 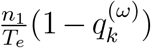 where 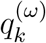 is the probability that the strain goes extinct during the epidemic burnout phase. Established strains are lost through two processes at two timescales: 1) burnout during new strain invasions, with probability 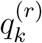 over the interval *T*_*e*_, 2) Long-term stochastic extinction, over a time window *E*(*T*_*Q*_), with probability 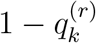. The combined death rate of established strains is a harmonic mean of the rates of these two processes.

#### ODE System for Strain Dynamics

We capture these dynamics with the following ODEs:

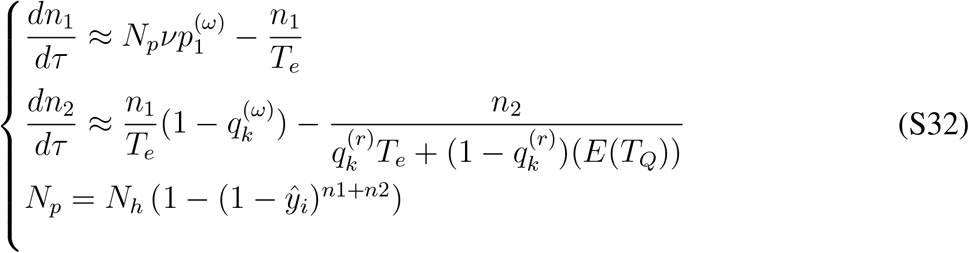

Transient strains (*n*_1_) contribute to established strains (*n*_2_) only if they survive the initial epidemic. The total number of strains at equilibrium is solved by setting 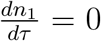 and 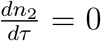. As described in the previous sections, the fizzle probability 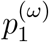, burnout probabilities 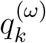 and 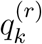, as well as the epidemic duration *T*_*e*_ and the long-term lifespan *E*(*T*_*Q*_) are dependent on the real-time number of strains, so they have to be recalculated given changes in *n*. With these quantities updated given a new set of (*n*_1_, *n*_2_), we will be able to numerically solve for the equilibrium status 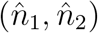.

#### Numerical solutions

We use the `nleqslv` function in the `R` package `nleqslv` (*110*) to solve for the equilibrium values 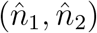, applying the Broyden method (*111*) and trust-region strategies (*112*). Providing the initial conditions of a guess on (*n*_1_, *n*_2_), together with the parameter array (*N*_*h*_, *R*_0_, *ε, ε*_1_, *ν, ρ*), we aim to solve for the pair 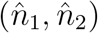 at the stable equilibrium, with where both eigenvalues from the Jacobian matrix have negative real parts. To ensure stability and accuracy, we test multiple initial conditions and accept solutions with the minimum sum of absolute values 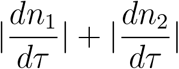, at least satisfying 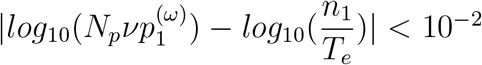 and 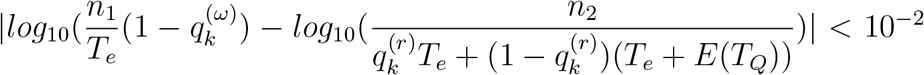. We prioritize solutions where the Jacobian matrix has negative eigenvalues (indicating local stability). If no stable solution is found, we return the one with the minimum residual 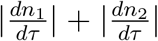. If no solution meets the tolerance criteria, we conclude that no strains can persist, and set *n* = 0. The code for MultiSEED is available at GitHub: https://github.itap.purdue.edu/HeLab/MultiSEED.

### 1.6 Stochastic compartmental model of strain dynamics using Markov chain methods

In addition to deriving the equilibrium value of 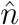 using Eq. S32, we also formulate a stochastic version of the model to simulate the temporal trajectories of strain dynamics using a continuous-time Markov process. This approach is based on four types of events derived from the deterministic equation: the birth and death of both transient (*n*_1_) and established (*n*_2_) strains. The rates of these events are as follows: emergence of new transient strains: 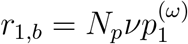, burnout of transient strains: 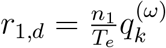, transition of transient to established strains: 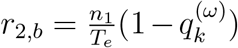, and the extinction of established strains: 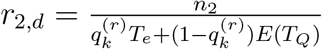. Given an initial state defined by *n*_1_ and *n*_2_, the Markov process proceeds by generating events based on these rates. The time to the next event is sampled from an exponential distribution with rate *r*_*t*_ = *r*_1,*b*_ + *r*_1,*d*_ + *r*_2,*b*_ + *r*_2,*d*_. The type of event is then selected probabilistically, weighted by its relative rate. After each event, the state variables *n*_1_ and *n*_2_ are updated accordingly, along with the simulation clock. The process terminates either when a predefined time limit is reached or when all strains go extinct (i.e., *n*_1_ = *n*_2_ = 0). For the Markov simulations shown in Figs. 2–4 of the main text, we initialized the system with one established strain (*n*_1_ = 0, *n*_2_ = 1) and ran 1,000 independent simulations for each parameter set to estimate the expected strain diversity over the specified co-evolutionary time period. This Markov-based stochastic compartmental model can similarly be used to track the evolution of strain diversity over time from arbitrary initial conditions.

### 1.7 Agent-based model of multi-strain SIRS dynamics

We developed an agent-based stochastic model (ABM) to simulate the eco-evolutionary dynamics of multi-strain transmissions (GitHub: https://github.itap.purdue.edu/HeLab/MultiSEED). Results from the stochastic simulations were used to verify our numerical approximations described in previous sections. The model tracks each infection within each host explicitly. Each infection is associated with a specific strain. Multiple infections of heterogeneous strains within the same host are allowed, and the transmissibility (*β*_*e*_) per infection will be reduced to 1*/m* if the host has *m* concurrent infections. After recovering from an infection by strain *i* (1*/γ*), the host gains specific immunity against strain *i*. This specific immune memory is lost at the rate of *δ*. Hosts who have been infected or have cleared infections gain a general protection against the disease, acquiring a reduced susceptibility towards future infections from heterogeneous strains (*ρ*). The rate of newborns in the population and the death rates of hosts are kept constant (*µ*). Infections do not cause host deaths. New strains are innovated from existing infections during their generation time, which includes the incubation period and infectious period. Note that *ν* is in the unit of *τ* = *t/*(*γ* + *µ*), so the strain innovation rate in the simulation is scaled to be *ν* * (*γ* + *µ*) per infected host. The copy number of each strain is tracked throughout the simulation. Origin and death times of strains are also recorded. Simulations are initiated with *N*_*h*_ naïve hosts, characterized by an exponentially distributed age profile with a mean life expectancy of 1*/µ*. When a host reaches its life expectancy and is removed from the simulation, a new naïve host is born. Strain frequencies, host prevalence, and immune profiles are sampled every 30 days from the transmission dynamics. Simulations are stopped when host prevalence and strain diversity have reached a steady state. Simulations may also terminate if the disease is cleared and there remain no infections in the system.

The ABM was developed to be as close as possible to the equation-based multi-strain model. However, the age structure of disease prevalence and immune profile, which naturally emerges from the ABM, is not accounted for in the ODE systems. The discrepancy will generate slight differences between the results from ABM and the numerical methods described above.

### 1.8 Empirical ranges of parameters in common infectious diseases

In this section, we present empirical estimates for the key parameters used in our model for several well-studied infectious diseases. The host population size *N*_*h*_ is estimated based on typical host behavior and the transmission scale of each pathogen. For each disease, we estimate values from the literature for the following parameters: the strain innovation rate (*ν*), cross-immunity susceptibility (*ρ*), the basic reproduction number (*R*_0_), and parameters used to calculate the relative resource replenishment rate *ε*. As introduced in Section 1.1, we define 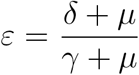, where *δ* is the immunity loss rate, *γ* is the recovery rate, and *µ* = 0.02 year^−1^ is the host birth/death rate, assuming an average human lifespan of 50 years. When estimating *γ*, we account for the full generation time of the infection, which includes both incubation and infectious periods, as consistent with generation interval of the SEIR model framework (see Eq. 3.5 in (*113*)). For the strain innovation rate *ν*, we estimate it by calculating the observed genetic divergence between known strains and dividing by the expected time to accumulate such divergence, based on known mutation or recombination rates. Lastly, we investigate the origin time of the disease to estimate the length of its co-evolutionary history with the host. This period is critical for determining whether the currently observed strain diversity reflects a long-term evolutionary equilibrium.

#### 1.8.1 Influenza A

Influenza A is highly transmissible in large populations. It has been suggested that large host populations enable the antigenic drift in influenza A and thus the lineage turnover (*55*). Therefore, We assume a relatively large host population size of *N*_*h*_ = 10^6^. The basic reproduction number *R*_0_ of influenza A is often estimated to be between 1.2 and 1.85, excluding the estimates for the initial 1918 epidemic and in confined settings, where extreme values were reported (*62*). To estimate the strain innovation rate, we focus on the HA1 receptor-binding domain, which consists of 329 amino acids (*56*). The per-nucleotide mutation rate is approximately 2 × 10^−5^ per cellular infection (*57*), with roughly two viral replication cycles per day (*58*), yielding a daily rate of 4 × 10^−5^. Given that the incubation and infectious periods each span 2–4 days (*59–61*), we estimate a generation time of 7 days, resulting in a per-generation mutation rate of 4 × 10^−5^ × 7 per nucleotide. Since strains with deleterious mutations are subject to strong purifying selection, the strain innovation speed should be scaled by the fraction of beneficial mutations, which is estimated to be 5*/*57 = 0.088 according to (*57*). The mean pairwise amino acid divergence among influenza A strains is about 13 residues (*38*), suggesting that a new strain requires a 3.95% change in amino acid sequence (13*/*329). Therefore, the strain innovation rate is 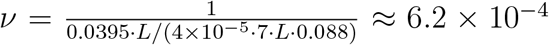, where *L* is the nucleotide length of the HA1 domain. We thus take the magnitude of 10^−4^ as the strain innovation rates per infection lifespan. We use the half-life of immune memory to calculate the strain-specific immune loss rate, assuming immunity over time follows an exponential decay. Protection against influenza A strains wanes by half in 3.5 to 7 years (*26*), which represents the median duration of immune memory. Therefore, we parameterize the strain-specific immunity to last from 3.5 to 7 years. Given these values, the replenishment ratio *ε* ranges from approximately 0.003 to 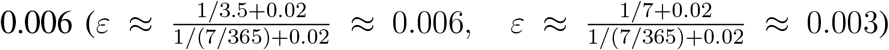. For cross-immunity, vaccine efficacy studies report that both live-attenuated and trivalent inactivated vaccines offer approximately 50% protection against mismatched strains (*63*). We therefore assume cross-susceptibility *ρ* = 0.5, indicating that prior infection reduces susceptibility to heterologous strains by half. Finally, the most recent common ancestor of influenza A hemagglutinin dates to around 1056 AD (*64*), suggesting that the disease has been co-evolving with humans for around 1000 years.

#### 1.8.2 Pneumococcus (*Streptococcus pneumoniae*)

The second example is the bacterial pathogen *Streptococcus pneumoniae*, which has more than 90 different serotypes detected based on the composition of its capsule polysaccharide (*6*). Since bacteria can be easily transmitted in large urban populations at the magnitude of one million (*79*), we set *N*_*h*_ = 10^6^. The basic reproduction number of *Streptococcus pneumoniae* has mostly been estimated to be no greater than 2 (*89, 90, 114*). In one specific study covering daycare in multiple countries, the basic reproduction number varies from 1.23 to 4.77 (*91*). However, the focal populations only include young children, and the ones with *R*_0_ *>* 2 have quite small sample sizes, and the only population with sample size larger than 1000 has *R*_0_ = 1.23. Therefore, we assume the range of *R*_0_ for this bacterial pathogen is 1.23 − 2. The carriers of this bacteria in their nose can spread the infection without showing symptoms, and the carriage duration may vary from 6 weeks to almost 5 months, according to one study (*80*), with a median of 132 days reported in another study (*81*). Thus, we assume that the generation period is approximately 120 days, equivalent to three generations of infections per year. For pneumococcus, the main source of genetic changes is recombination rather than spontaneous point mutations (*115, 116*), we will use the recombination rates to estimate *ν*. The capsule polysaccharide locus (cps), whose genotypes define serotypes, recombines at a rate of approximately 0.005 per year (*82*). The length of cps is 10-30 kilobases (*83*), so that the recombination rate is 0.005*/*20000 = 2.5 × 10^−7^ per base per year when assuming the length is 20kb. Accordingly, the number of recombination events in a generation period of 120 days over the antigen coding region is 2.5 × 10^−7^ × *L/*365 × 120, where *L* is the length of the antigen coding region. The genetic difference between cps serotypes can be as small as 3% (*84, 85*). Therefore, the number of events required for forming a new strain is around 0.03 × *L*, suggesting that 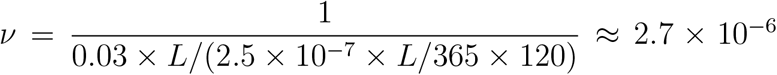. Thus, we use 10^−6^ as the magnitude of strain innovation rate for pneumococcus. Serotype-specific immune memory wanes with age, and older individuals are a high-risk group for reinfection; however, they still retain certain levels of B cells, sometimes even into their 100s (*86, 87*). Therefore, the lower bound of *ε* is estimated as 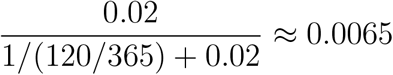 assuming lifelong immunity. Another study indicates that the IgG antibody level can drop from 78% to 58% within 6 years after vaccination in elderly individuals (*88*). If we assume an exponential decay of immune memory over these 6 years (18 generations as there are 3 generations per year), we can calculate the decay rate by solving the equation 58 = 78 × *e*^−18*λ*^. Since *λ* ≈ 0.016, the mean lifetime of antibodies can be approximated as 1*/λ* ≈ 61 generations in this scenario, which is roughly 20 years. Thus, we estimate the upper bound of *ε* assuming the waning immunity over 20 year, so that 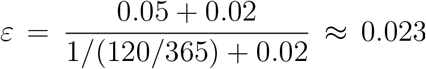. Studies on different conjugate vaccines have commonly indicated that the efficacy against all pneumococcal pneumonia is around 30% (*92–94*),while serotype-independent protection in children is around 40% (*95*). These results suggest that we can roughly estimate that *ρ* = 0.7. The common ancestors of *Streptococcus pneumoniae* and its close relatives date back as far as a million years ago (*85*). It had a tiny effective population size until the recent expansion of the human population in the last 120,000 years. Therefore, we assume that the circulating time of *Streptococcus pneumoniae* in human populations is around 100,000 years.

#### 1.8.3 HIV-1

HIV-1 is transmitted through body fluids, indicating that its spread is highly dependent on individual host behaviors and is typically constrained to small regional populations. Such diseases can transmit within a population with around 10,000 hosts (*65*). Accordingly, we set the host population size to *N*_*h*_ = 10^4^. The infectious period of HIV-1 is approximately one year (*66*). Both the recombination and mutation rates of HIV-1 fall within the range of 1.4 × 10^−5^ to 2 × 10^−4^ per nucleotide per viral replication cycle (*67–70*). Given the short half-life of infected cells (i.e., about two days (*71*)), we estimate that the infection undergoes approximately 90 replication cycles per year. The key antigenic region of the HIV-1 genome is the Gag domain, which spans roughly 1500 nucleotides. One study suggests that 5% of HIV-1 mutations are beneficial (*72*), while another reports that subtypes differ by 17% in amino acid sequence within the Gag region (*73*). Assuming that 70-75% single-nucleotide mutations are non-synonymous, 17% amino acid changes map to around 20% nucleotide divergence among strains. We estimate the strain innovation rate *ν* as follows: assuming 0.05 × 1 × 10^−4^ × 1500 ÷ 4 × 365 beneficial recombination events per generation and 0.2 × 1500 genetic changes are required to produce a new strain, 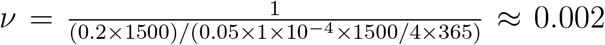. This places the magnitude of *ν* around 10^−3^ per generation period. Estimating the strain-specific immune loss rate *δ* is challenging. In untreated individuals, antibody levels tend to stabilize long-term (*74*), and superinfections typically occur only when encountering genetically distinct strains (*75*). Thus, we assume *δ* ≈ 0, representing nearly lifelong strain-specific immunity (*76*). Using this assumption, we estimate the bounds of the replenishment rate *ε* under different host mortality rates: for a 75-year lifespan 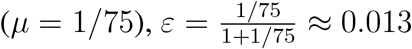, whereas for a 25-year lifespan 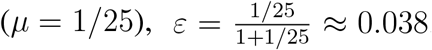. The basic reproduction number *R*_0_ for HIV-1 is estimated between 1 and 3, with a 95% HPD interval of [1.6,3.1] when assuming general epidemiological parameters in ten subepidemics from one study (*77*). Regarding cross-immunity, a study reports a hazard ratio of 0.47 for superinfection compared to initial infections (*78*), suggesting partial protection. We therefore assume *ρ* = 0.5. Regarding the co-evolutionary history, HIV-1 has been present in humans for roughly 100 years. The earliest known human case was identified in 1959, while the most recent common ancestor of the HIV-1 M group is estimated to date back to around 1930 (*39, 40*).

#### 1.8.4 Malaria

Since falciparum malaria transmission is mostly endemic, and we set the resource host population size *N*_*h*_ to 10^5^, according to the reported population size in small to intermediate cities in Africa (*96*). The generation period of malaria varies significantly in treated and untreated patients, with a recovery time range of approximately 20 to 200 days (*97–99*). Here, we use 150 days, based on the duration of infection in naïve hosts. The antigenic diversity of *var* genes in the malaria parasite is largely generated through ectopic recombination (*117*); therefore, we use the recombination rate (around 1.8 × 10^−7^ per gene per day (*16*)) to estimate the innovation rate of new strains. The pairwise type sharing of *var* repertoires between malaria parasites tends to be kept at very low levels (mostly below 20%) to favor the persistence of parasites (*100*); thus, we assume that completely different *var* gene repertoires are required to form a new strain. In one generation of infection, assuming there are 60 *var* genes on one strain (*117*), we expect *n*_*r*_ = 1.8 × 10^−7^ × 150 × 60 recombination events, and the number of generations for recombination to occur on all genes is 60*/n*_*r*_. The strain innovation rate is calculated as follows, 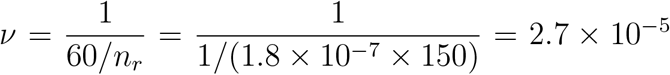 per generation. Thus, the magnitude of *ν* for malaria is around 10^−5^. The immune memory to one antigen can last for at least 3 years, and sometimes more than 8 years or even 16 years (*23,101–103*). We assume that the immunity to a strain will be lost to enable another infection by itself when the relatively long-lasting immune memory of antigens is lost, so here if we have *δ* = 1*/*8 then 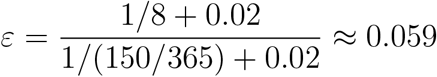 and *ε* ≈ 0.0336 when *δ* = 1*/*16. The basic reproduction number for malaria is difficult to estimate, as it varies widely from 1 to 3000 (*104*), because malaria is a vector-borne disease, and transmissibility depends on the host-vector behavior and density ratio. For cross-immunity among strains, the vaccine efficacy study on malaria strains indicated that the efficacy against unmatched strains is about 0.334 (*105*), suggesting that *ρ* ≈ 0.7. The malaria parasite *Plasmodium falciparum* was transmitted to humans from apes, and genetic diversity has accumulated within the past 10,000 years (*106, 107*). Here, we use 10,000 years as the co-evolutionary length between malaria parasites and humans.

## 2 Supplemental Figures

## 3 Supplemental Text

### 3.1 Negligible effects of the ratio of *ε*_1_ **to** *ε*

In our main text, when we explore the determinants of strain diversity, we regard *ε* as a single parameter, rather than studying its two components, *ε*_1_ and *ε*_2_, separately. However, as indicated in the ODE systems S7 and S9, sometimes *ε*_1_ and *ε*_2_ are involved separately in the compartmental model. Therefore, when *ε* = *ε*_1_ + *ε*_2_ is constant, it is possible that the relative difference between *ε*_1_ and *ε*_2_ can have some influence on the strain diversity, especially when general protection from cross-immunity is considered (*ε*_2_ is in equation set S9 but not in S7). However, such influence is usually negligible, as we will show here in this section.

When the ratio varies from 0.1 to 0.9, the effect of this ratio is generally negligible when the values of *ε* and *R*_0_ are intermediate or high (Fig. S9). This ratio mainly affects strain diversity under small values of *ε* or *R*_0_ near the persistence threshold. The lower this ratio is, which represents a higher replenishment from host births, the smaller threshold value of *ε* or *R*_0_ is required for the accumulation of strain diversity. This is because the replenishment of susceptible hosts (*ε*_2_) is more beneficial for the survival of new pathogen strains as it supplies naïve hosts, compared to the waning strain-specific immunity (*ε*_1_) that supplies hosts with protection from generalized immunity. Overall, the differences in strain diversity under different *ε*_1_*/ε* ratios are generally minor and less than one magnitude. Therefore, we simplify *ε*_1_ and *ε*_2_ as one parameter *ε* for factor exploration in the main text.

### 3.2 The number of possible equilibria

Sometimes, multiple equilibria can exist in the transmission system with positive feedback between genetic diversity and force of infection (*118*). When we obtain numerical approximations of strain diversity using MultiSEED, we observe scenarios with two possible equilibrium states, where the alternative equilibrium often comes with 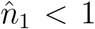 and 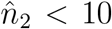, which probably represent the unstable equilibrium state that occurs when strains are first accumulating. We systematically examined the parameter space to identify regions where multiple equilibria may be found. We focus on the boundary region of persistence threshold set by *R*_0_ and *ε*, and fixed the combinations of *N*_*h*_, *ν* and *ρ* for the four focal human diseases. We use several initial conditions of different magnitudes in finding the equilibrium values of 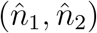. As Fig. S10 shows, multiple equilibrium states exist close to the persistence threshold with low *ε*. These low equilibria often have one or both values of *n*_1_ and *n*_2_ less than 1, and sometimes one to two magnitudes smaller than those of the corresponding high equilibria. For influenza A and HIV-1, which have a regime of strain replacement, the existence of two equilibria is relatively frequent over their empirical parameter space, particularly when the strains start to accumulate. For pneumococcus and malaria in regimes of coexistence, the existence of two equilibria is rare over their empirical parameter space, validating the stable co-circulation of multiple strains in such diseases. We then test the stability of these equilibria by evaluating the eigenvalues of the Jacobian matrices in finding 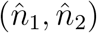. The eigenvalues from the numerical solver, `nleqslv`, can be affected by the initial conditions and are thus not 100% reliable. Nevertheless, we still find that the low equilibria are more frequently unstable compared to the high equilibria.

**Figure S1:**
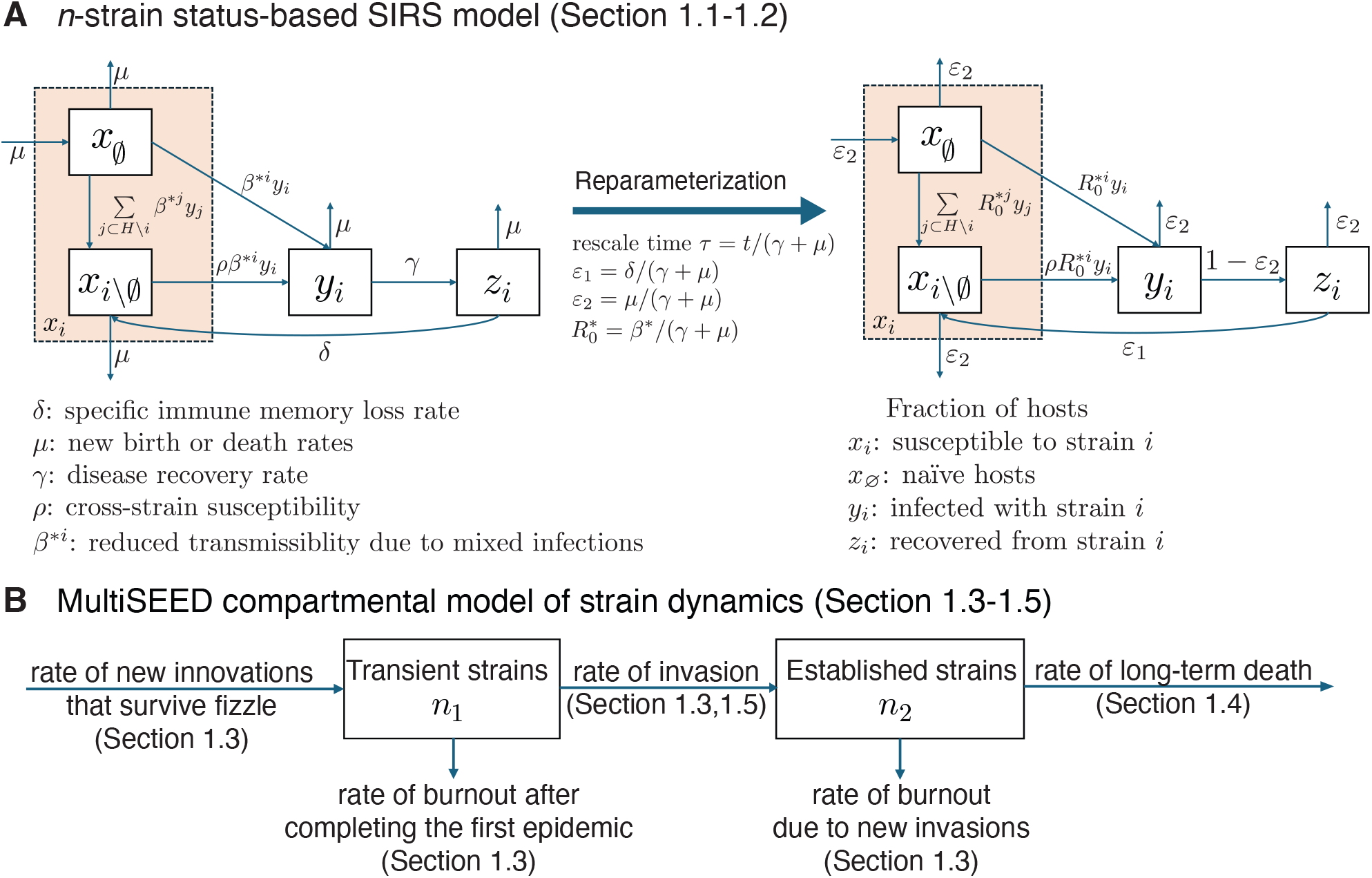
MultiSEED framework. **(A)** The deterministic *n*-strain status-based SIRS model. The definitions of host classes per strain and parameters are listed below. The fraction of susceptible hosts to strain *i, x*_*i*_, consists of *x*_∅_, which are complete naïve hosts, and *x*_*i\*∅_, which have past infection history. Fractions of hosts infected by strain *i* and recovered from strain *I* are respectively *y*_*i*_ and *z*_*i*_. Here, we assume that one host can be infected by multiple distinct strains, such that each strain will have reduced transmissibility *β*^*^. Disease recovery rate per strain is constant at rate *γ*. Previous infections will lead to general cross-immunity, resulting in reduced susceptibility (*ρ*). One specific strain *i* can infect hosts with no infection history, hosts infected by other strains, and hosts that lose strain-specific immune memory (with the waning rate *δ*). Birth or death rates of hosts are set to be constant *µ*. **(B)** The compartmental model of strain dynamics. We categorize strains into transient (*n*_1_) and established (*n*_2_) groups, tracking the rates of fizzle and burnout for transient strains, as well as the rates of burnout and long-term death for established strains.

**Figure S2:**
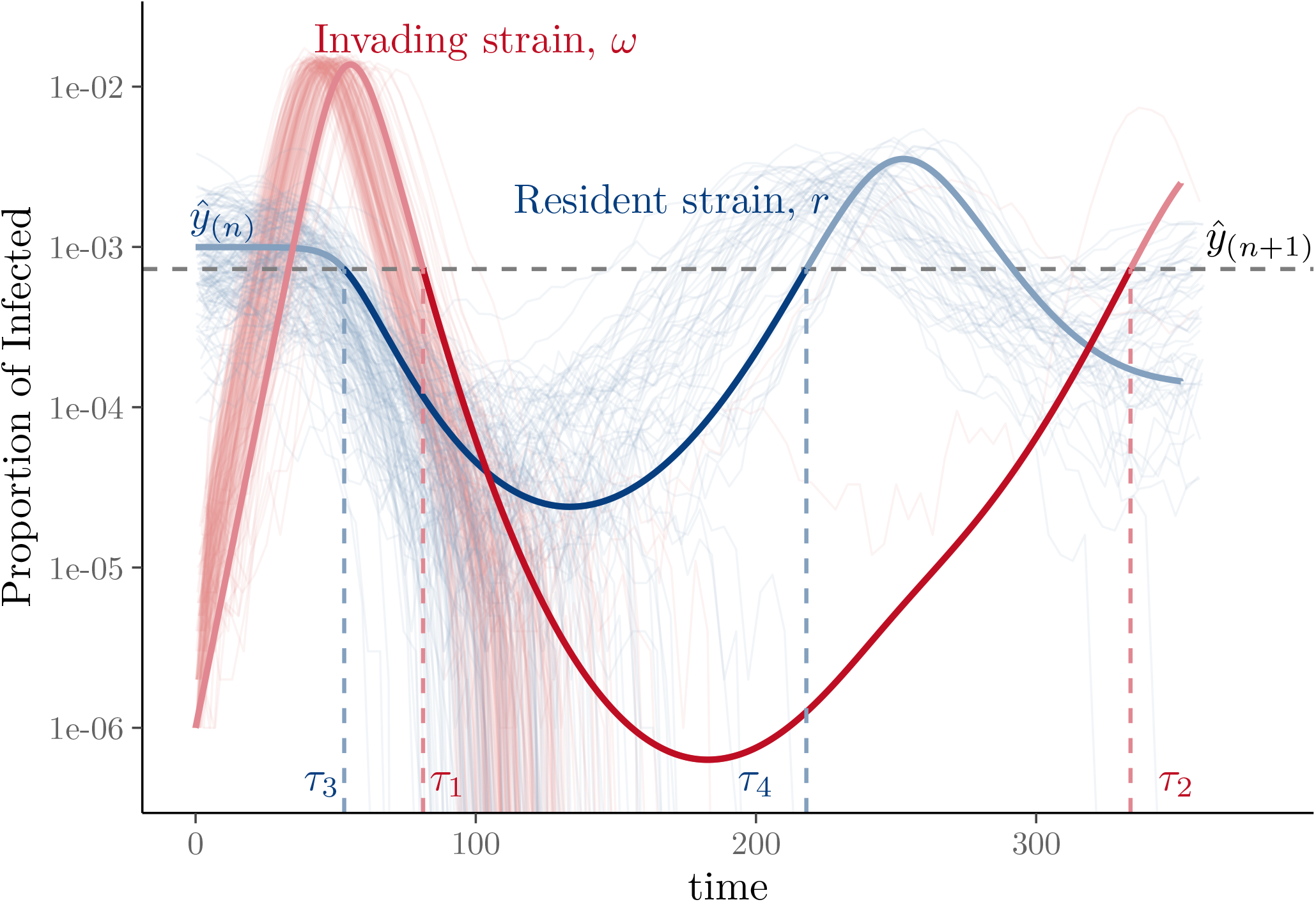
Illustration of deterministic and stochastic dynamics during a new strain invasion. Thick lines show the deterministic trajectories of the proportion of hosts infected with the invading strain *ω* (red) and a resident strain *r* (blue), obtained by numerically solving Eq.S10, assuming one resident strain (*n* = 1), a host population size of *N*_*h*_ = 10^6^, with parameters *ρ* = 0.5, *ε*_1_ = 0.0045, and *ε*_2_ = 0.0005. To capture stochastic effects, 1,000 agent-based simulations were run using the same parameter set, modeling the introduction of a single mutant strain into a system at endemic equilibrium (see Section 1.7 for model details). Translucent lines represent the 162 stochastic trajectories in which the invading strain survived the initial fizzle phase. Of these, only 2 strains successfully established, while 160 went extinct during the subsequent burnout phase. In addition, 112 resident strains were driven to extinction due to the epidemic triggered by the invading strain *ω*. These outcomes closely match the numerical approximations derived using the methods in Section 1.3: fizzle probability of the invading strain is 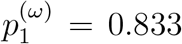; burnout probabilities of the invading strain and the resident strain are 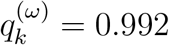 and 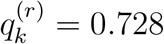, respectively.

**Figure S3:**
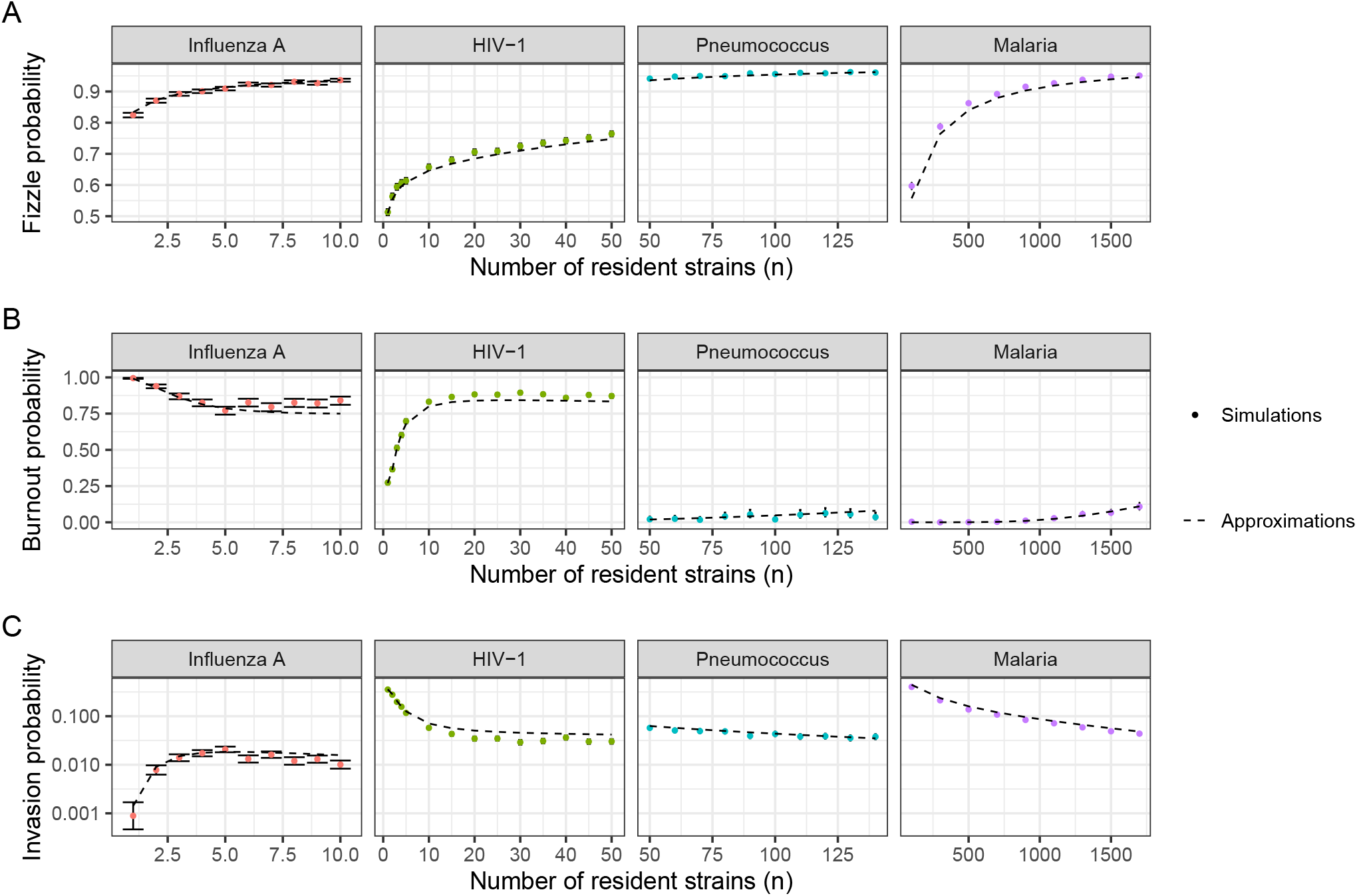
Comparison of predicted probabilities of stochastic processes at the ecological timescale between MultiSEED and agent-based stochastic simulations. **(A)** Fizzle probability, **(B)** burnout probability, and **(C)** invasion probability of a new strain are shown across four human diseases with different numbers of resident strains. Filled circles represent results from agent-based stochastic simulations, and the error bars represent 95% binomial proportion confidence interval. The parameter sets are identical to those described in Fig. 1D-E in the main text.

**Figure S4:**
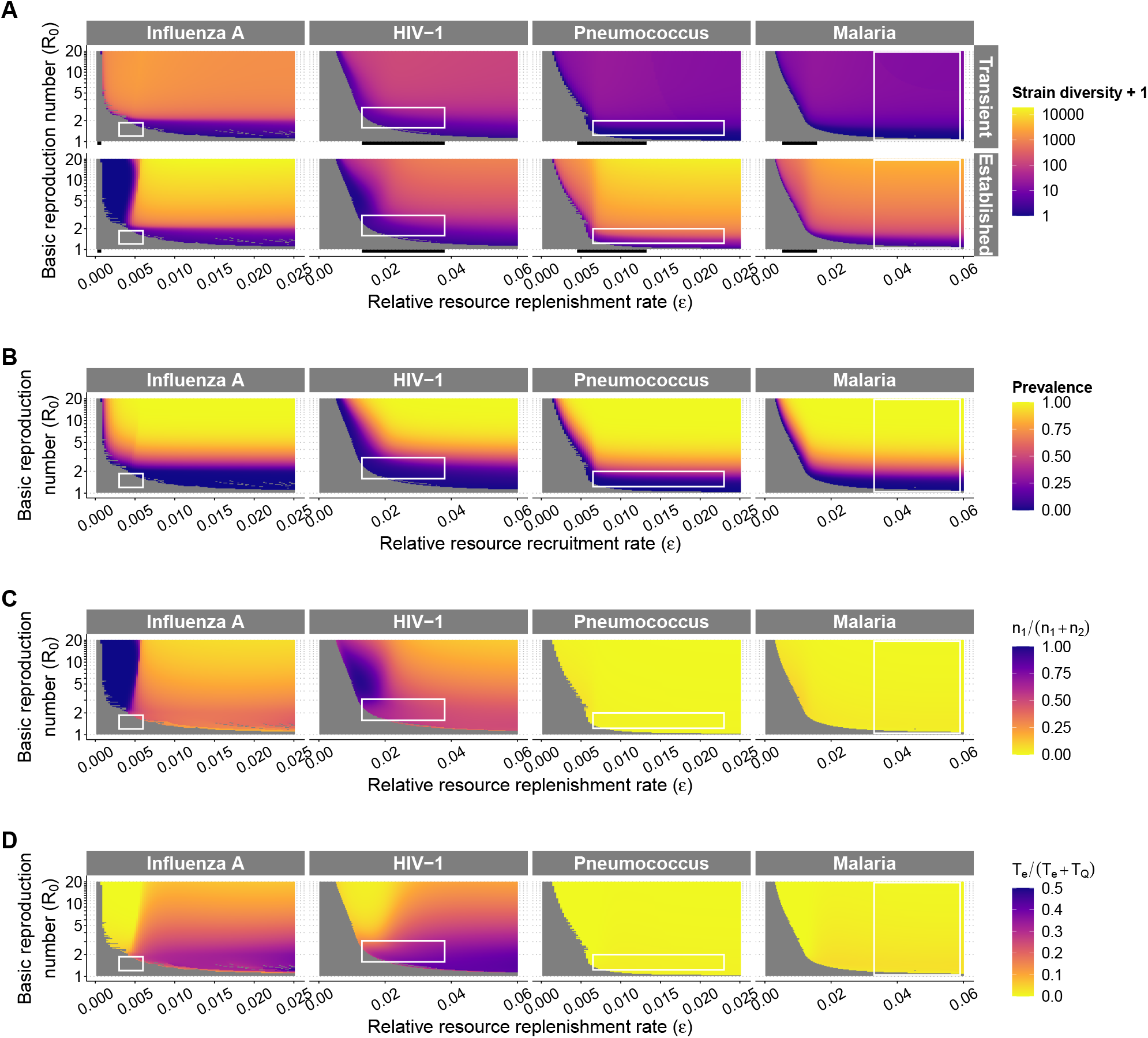
Predicted strain diversity, prevalence and the two quantities used in defining the replacement and coexistence regimes of the four focal diseases. The empirical ranges of *ε* and *R*_0_ are indicated by the white rectangles (see Table 1 in the main text for details). **(A)** Predicted numbers of transient and established strains at long-term equilibrium with varying *ε* and *R*_0_. Grid cells with zero strain diversity are shown in gray. The black bars below x axes indicate the resource replenishment rate through host birth with long-lasting strain-specific immunity (*ε* = *ε*_2_). **(B)** Prevalence. **(C)** The ratio of transient strains in all strains: *n*_1_*/*(*n*_1_+*n*_2_). **(D)** The ratio between the duration of the first epidemic and the expected extinction time at the evolutionary time scale, *T*_*e*_*/*(*T*_*e*_ + *T*_*Q*_). In Fig. 2C in the main text, the regime of coexistence is defined when both quantities are below 1/3 in **(C)** and **(D)** here. Whenever either quantity is no less than 1/3 outside of the extinction regime, we define the dynamics to be replacement.

**Figure S5:**
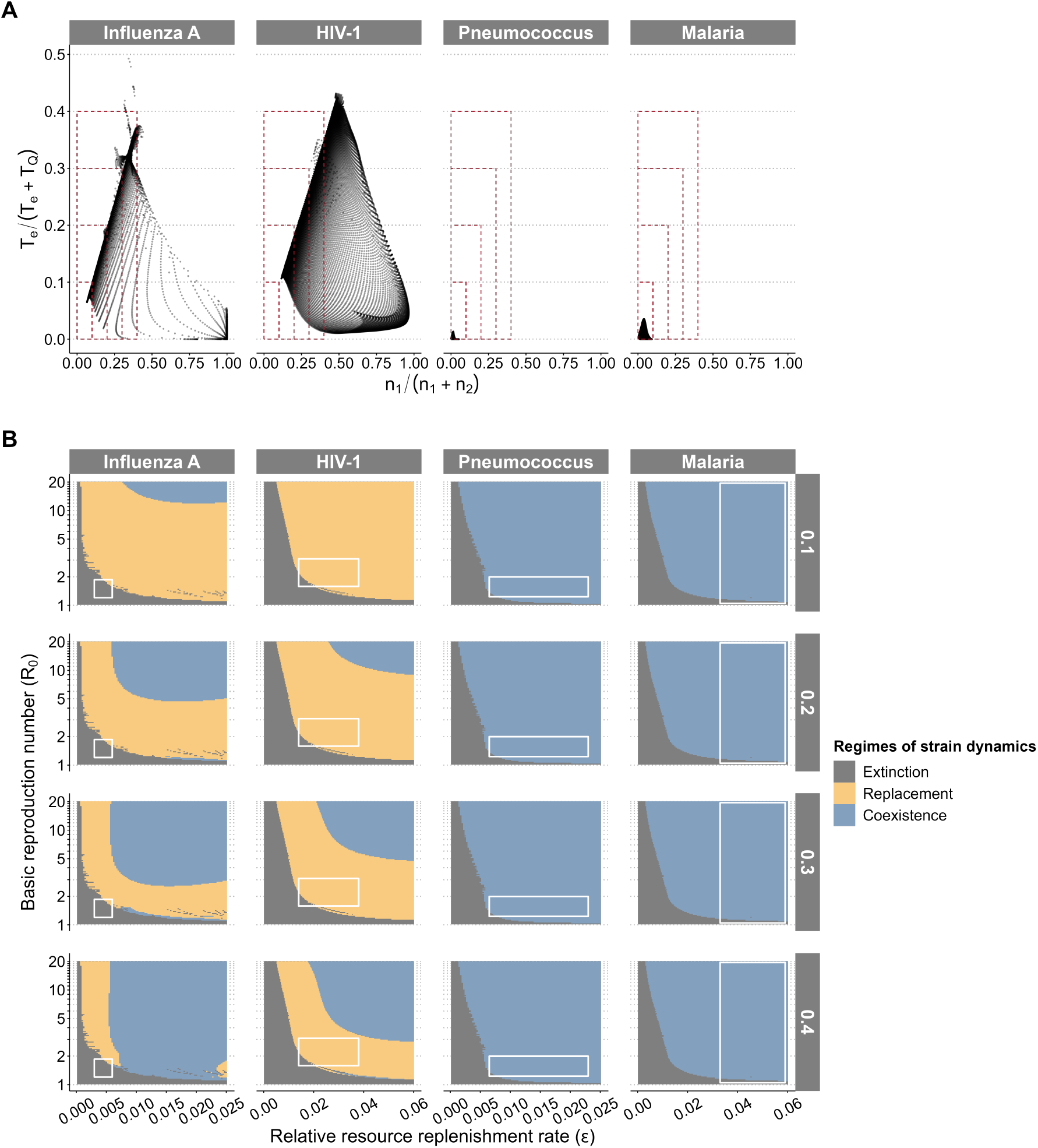
Effects of the cutoff values in regime delineation. **(A)** The correlation between the two quantities defining regimes of strain dynamics mentioned in Fig. S4: *n*_1_*/*(*n*_1_ + *n*_2_) and *T*_*e*_*/*(*T*_*e*_ + *T*_*Q*_). Red rectangles in dashed lines indicate the four cutoff values considered in defining the regime. Points within the rectangles are considered coexistence; otherwise, replacement occurs. For the broad parameter spaces of *N*_*h*_, *ν* and *ρ* in influenza A and HIV-1, the regimes are mostly replacement, but sometimes coexistence depending on the cutoff values. For the parameter spaces of pneumococcus and malaria, the regime is always coexistence, as points accumulate at the bottom-left corner with extremely small values of both quantities. **(B)** The regimes of strain dynamics when using different cutoff values of the two quantities. Coexistence is suggested when both quantities are below the cutoff value.

**Figure S6:**
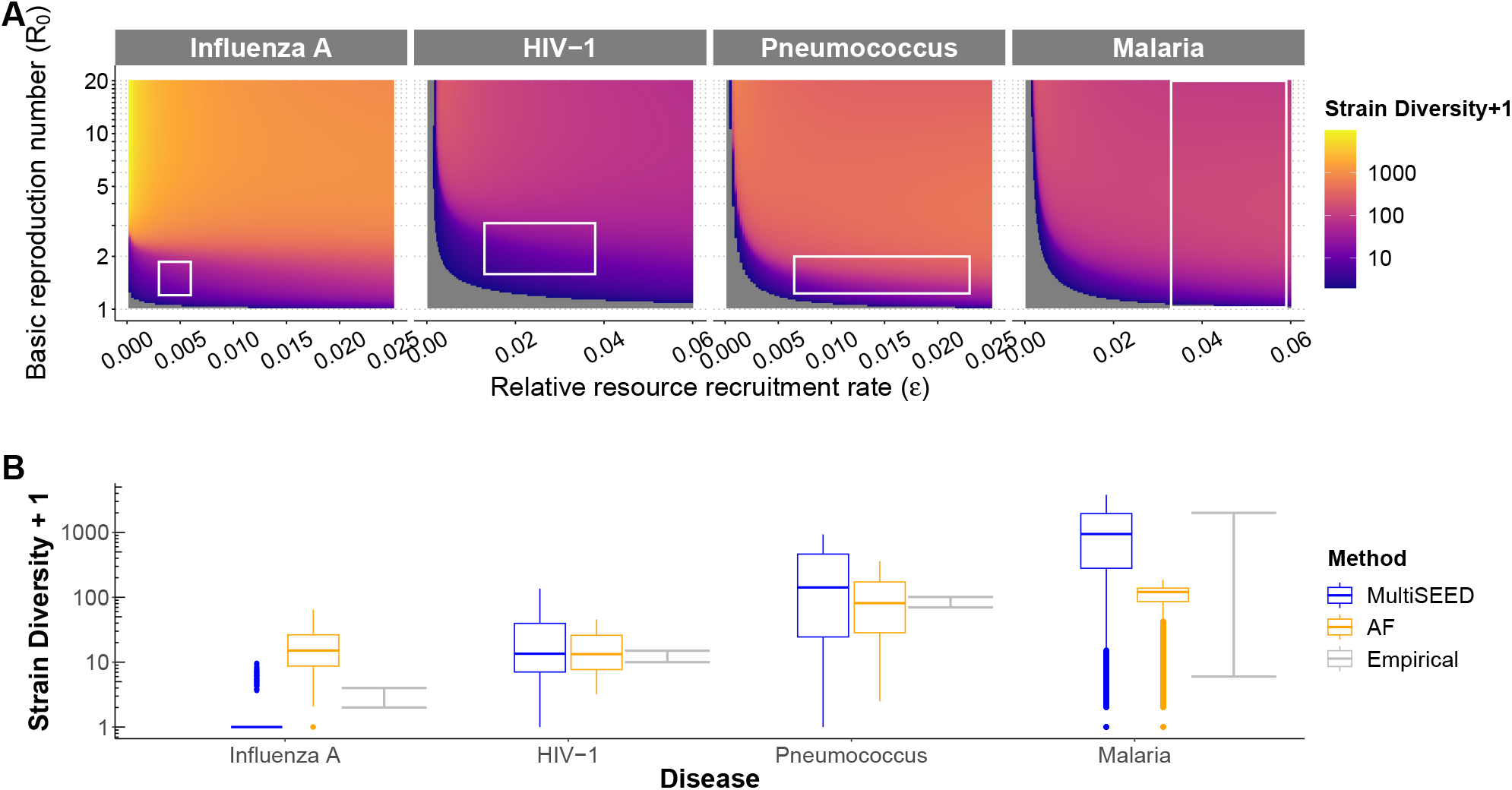
Comparison of strain diversity predicted by MultiSEED and by the methods from Abu-Raddad & Ferguson (*9*), here referred as “AF-model”. **(A)** Grids of strain diversity estimated using the AF-model considering strain innovation and long-term stochastic strain death as a function of *ε* and *R*_0_. In the original study of the AF-model, replenishment of pathogen resources occurs only through the births of susceptible hosts, equivalent to *ε*_2_ in MultiSEED, without considering waning immunity. To make the model outputs comparable, we supply *ε*_2_ in AF-model with our values of resource replenishment *ε*. **(B)** Comparison of strain diversity predicted by the AF-model and MultiSEED, together with the empirical ranges. Blue boxes indicate predicted strain diversity from MultiSEED (*n* = *n*_1_ + *n*_2_), and orange ones represent the predicted strain diversity from the AF-model. Grey bars stand for the observed range of strain diversity in these diseases.

**Figure S7:**
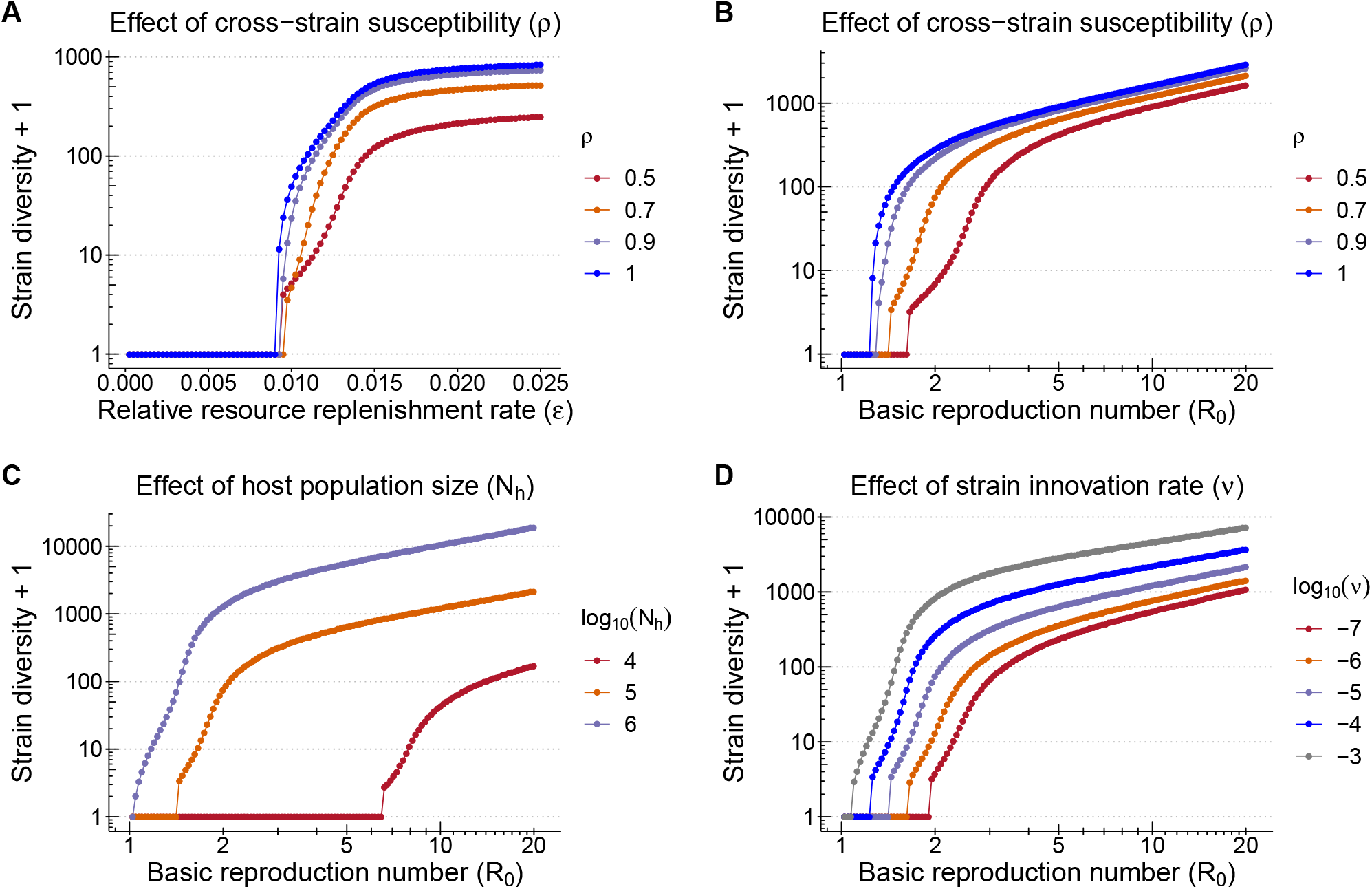
Effects of parameters on strain diversity and the persistence threshold along the *R*_0_ **or** *ε* axis, as a supplementary for Fig. 3B-C in the main text. **(A-B)** The effects of cross-strain susceptibility (*ρ*) along the *ε* and *R*_0_ axis. **(C-D)** The effects of host population size (*N*_*h*_) and strain innovation rate (*ν*) along the *R*_0_ axis. The default parameter values are *N*_*h*_ = 10^5^, *ν* = 10^−5^, *ρ* = 0.7, *ε* = 0.015 and *R*_0_ = 3. In each plot, one parameter on the x-axis and one parameter indicated by the legend are changed, while others are fixed.

**Figure S8:**
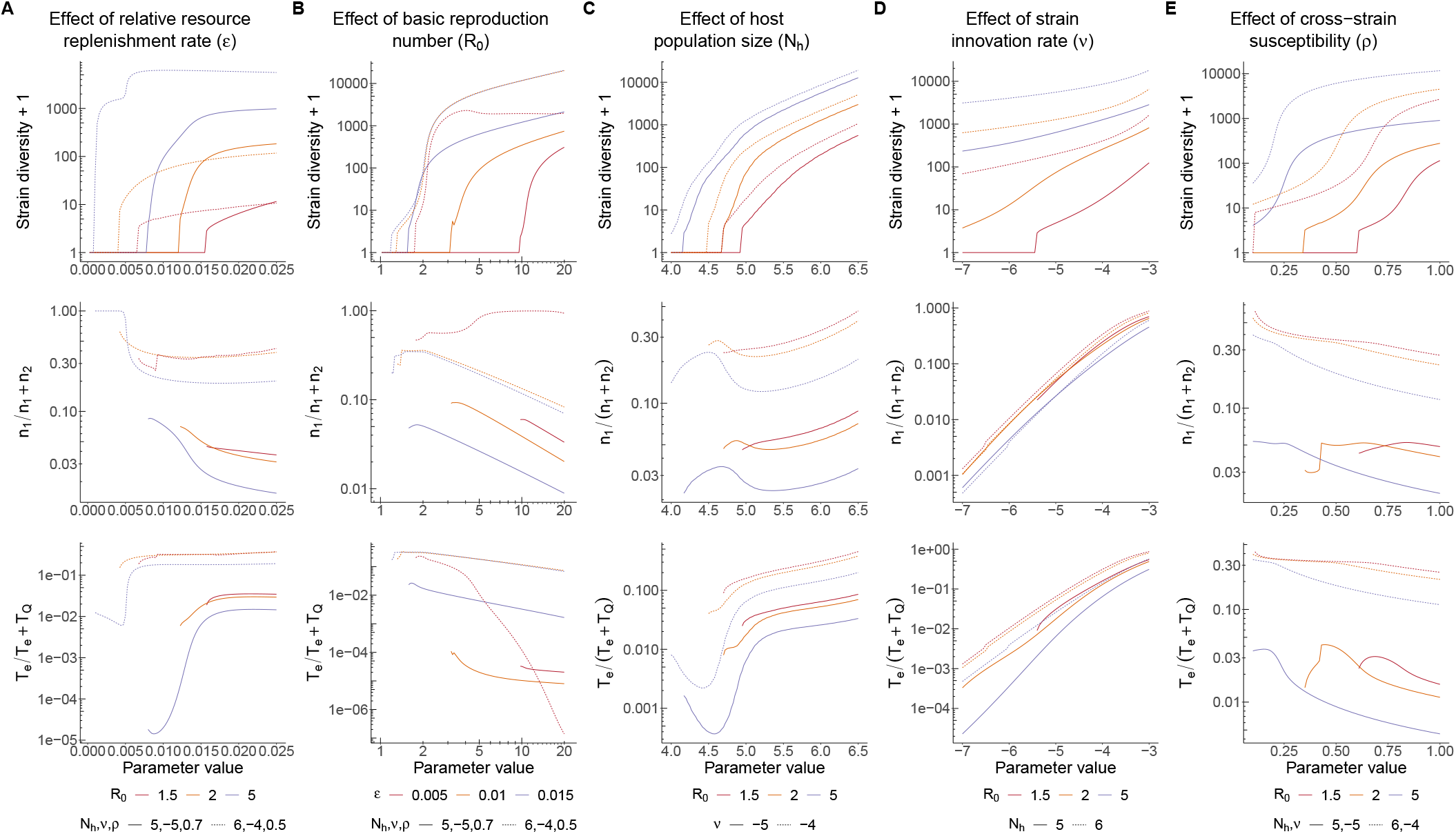
Effects of different parameters on long-term strain diversity and regimes of strain dynamics. Plot rows one, two and three correspond to the responses of strain diversity and the two criteria for classifying regimes of strain dynamics given varying parameters. Parameters *N*_*h*_, *ν* are always at *log*_10_ scale. **(A)** The effects of the relative resource replenishment rate *ε*. Two parameter combinations are tested here: *N*_*h*_ = 10^5^, *ν* = 10^−5^, *ρ* = 0.7, *ε*_1_*/ε* = 0.75, representing the broad parameter region of malaria, and *N*_*h*_ = 10^6^, *ν* = 10^−4^, *ρ* = 0.5, *ε*_1_*/ε* = 0.9, representing the broad parameter region of influenza A. **(B)** The effects of basic reproduction number *R*_0_. The same parameter combinations in **A** are used. Here *ε* = 0.003 is not included because for the parameter combination *N*_*h*_ = 10^5^, *ν* = 10^−5^, *ρ* = 0.7, strain diversity is always 0 when *ε* = 0.003. **(C)** The effects of host population size *N*_*h*_, with fixed parameter values *ε* = 0.015 and *ρ* = 0.7. **(D)** The effects of strain innovation rate *ν*, with *ε* = 0.015 and *ρ* = 0.7. **(E)** The effects of cross-strain susceptibility *ρ*, with *ε* = 0.015.

**Figure S9:**
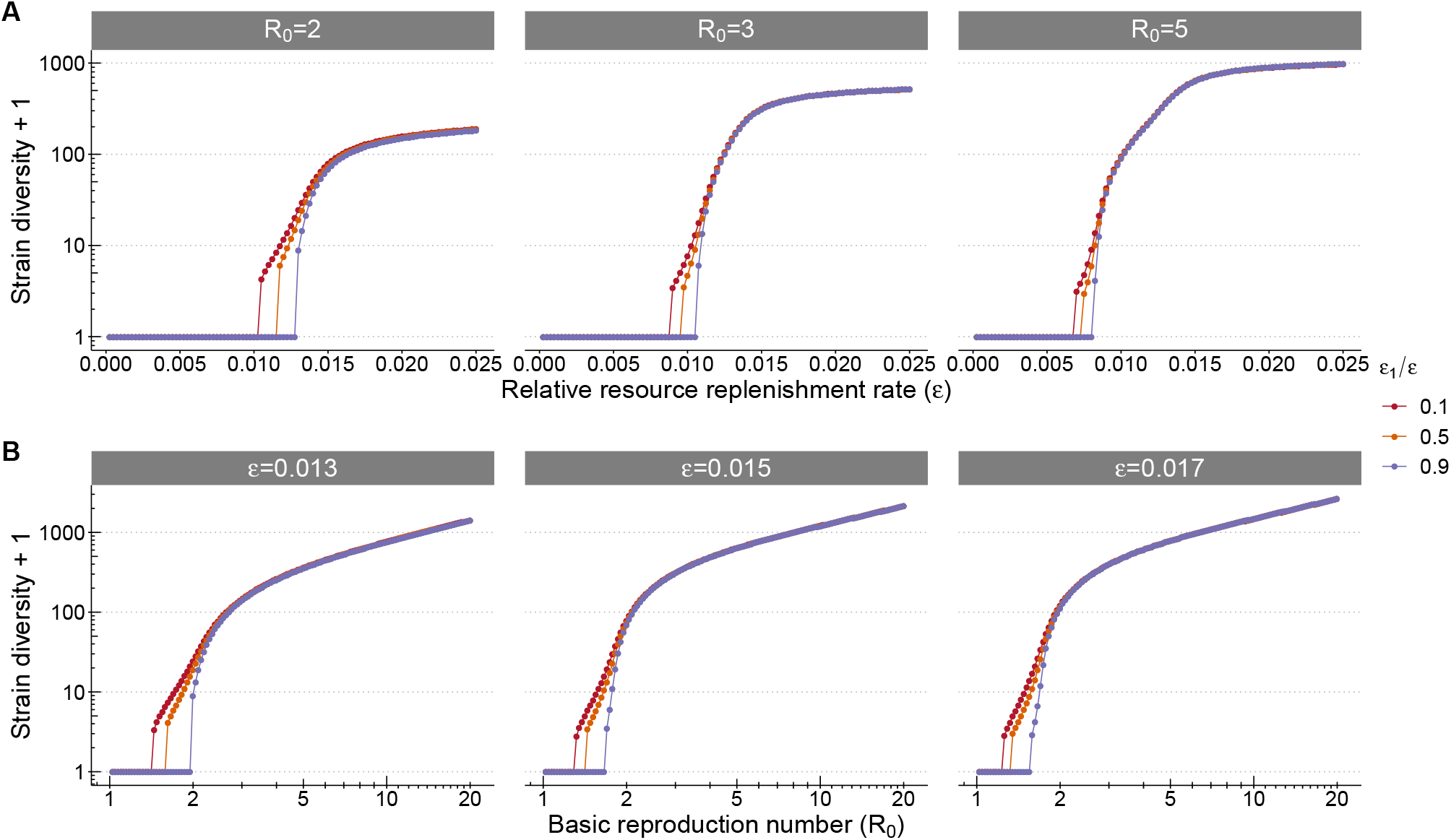
The effect of 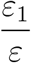 on long-term strain diversity, under different values of *R*_0_ and *ε*. The values of other parameters are *N*_*h*_ = 10^5^, *ν* = 10^−5^ and *ρ* = 0.7.

**Figure S10:**
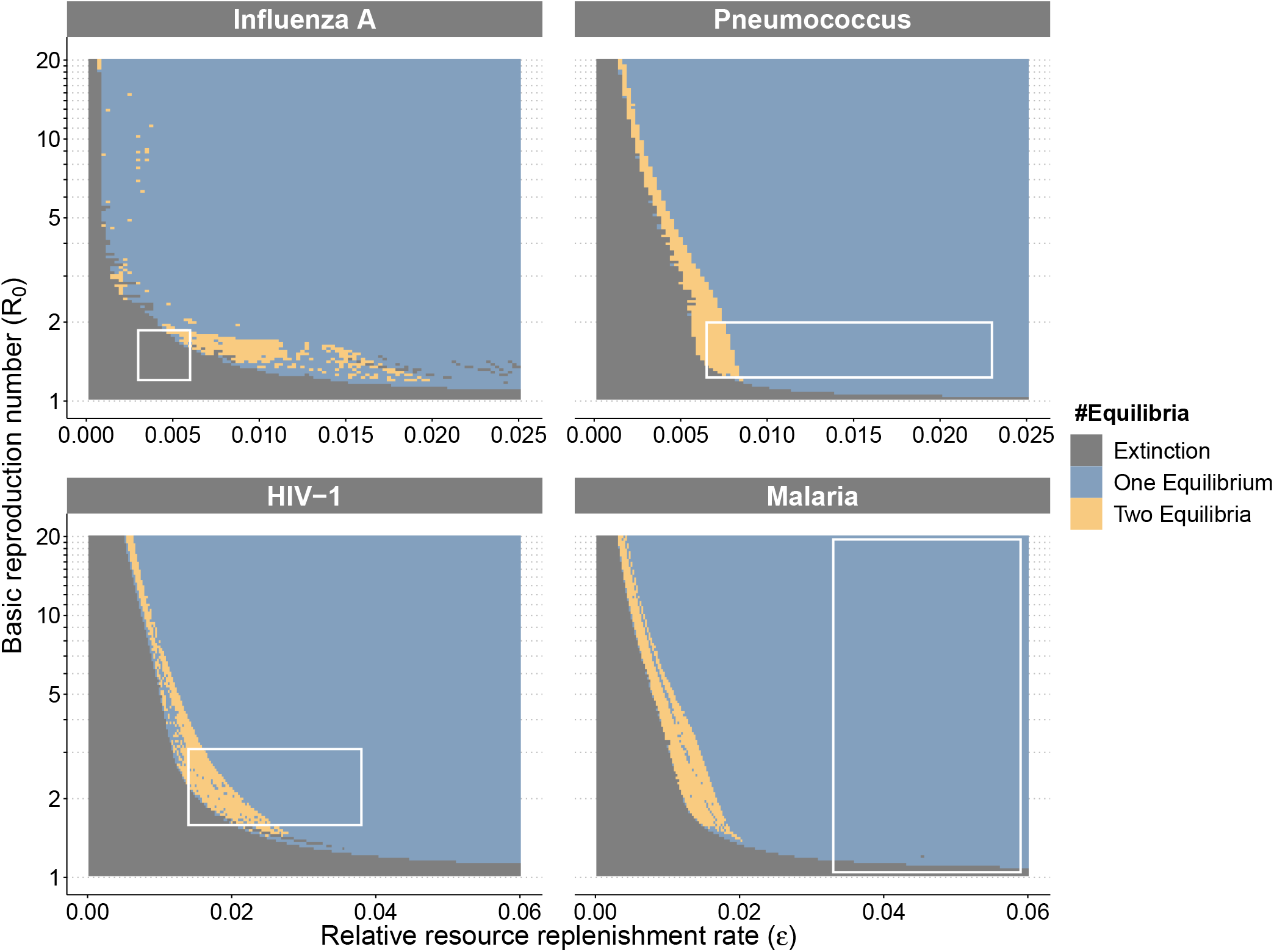
Estimated number of equilibria using MultiSEED. Each subplot shows the number of equilibrium under different combinations of *ε* and *R*_0_ for the four human diseases. The empirical ranges of *ε* and *R*_0_ are indicated by the white rectangles, identical to Figs. 3 and 4 in the main text. We delineate the boundary region to search for alternative equilibria as follows: For each value of *R*_0_, we first locate the smallest value of *ε* that leads to a positive strain diversity (*ε*_*s*_). Then we examine the range from *ε*_*s*_ to (*ε*_*s*_+0.01) or the upper bound of *ε*. To find the number of equilibria, we search for multiple initial conditions in each grid cell, and keep all the solutions found for 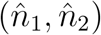, as described in Section 1.5. The solutions of equilibria are considered identical when the values of 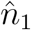 or 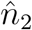 differ by less than 0.01 on the log scale.

**Table S1:** Transition rates for the stochastic reparameterized single-strain SIRS model: Eq. S5

| Event | Type of transition | Transition rate $g_j(x, y)$ |
| --- | --- | --- |
| New birth or immunity loss | $(S, I) \rightarrow (S + 1, I)$ | $N_h g_1(x, y) = \varepsilon_1(N_h - S - I) + \varepsilon_2 N_h$ |
| Death of susceptibles | $(S, I) \rightarrow (S - 1, I)$ | $N_h g_2(x, y) = \varepsilon_2 S$ |
| Death or recovery of infected | $(S, I) \rightarrow (S, I - 1)$ | $N_h g_3(x, y) = I$ |
| Infection of susceptibles | $(S, I) \rightarrow (S - 1, I + 1)$ | $N_h g_4(x, y) = R_0 I S / N_h$ |

